# Discovery of novel representatives of bilaterian neuropeptide families and reconstruction of neuropeptide precursor evolution in ophiuroid echinoderms

**DOI:** 10.1101/129783

**Authors:** Meet Zandawala, Ismail Moghul, Luis Alfonso Yañez Guerra, Jérôme Delroisse, Nikara Abylkassimova, Andrew F. Hugall, Timothy D. O’Hara, Maurice R. Elphick

**Author notes:** Contributed equally (Ismail Moghul, Luis Alfonso Yañez Guerra, Jérôme Delroisse). Correspondence to: Prof. M.R. Elphick and Dr. M. Zandawala, and. Luis Alfonso Yañez Guerra. Jérôme Delroisse.

## Abstract

Neuropeptides are a diverse class of intercellular signaling molecules that mediate neuronal regulation of many physiological and behavioural processes. Recent advances in genome/transcriptome sequencing are enabling identification of neuropeptide precursor proteins in species from a growing variety of animal taxa, providing new insights into the evolution of neuropeptide signaling. Here detailed analysis of transcriptome sequence data from three brittle star species, *Ophionotus victoriae, Amphiura filiformis* and *Ophiopsila aranea*, has enabled the first comprehensive identification of neuropeptide precursors in the class Ophiuroidea of the phylum Echinodermata. Representatives of over thirty bilaterian neuropeptide precursor families were identified, some of which occur as paralogs. Furthermore, homologs of endothelin/CCHamide, eclosion hormone, neuropeptide-F/Y and nucleobinin/nesfatin were discovered here in a deuterostome/echinoderm for the first time. The majority of ophiuroid neuropeptide precursors contain a single copy of a neuropeptide, but several precursors comprise multiple copies of identical or non-identical, but structurally-related, neuropeptides. Here we performed an unprecedented investigation of the evolution of neuropeptide copy-number over a period of ~270 million years by analysing sequence data from over fifty ophiuroid species, with reference to a robust phylogeny. Our analysis indicates that the composition of neuropeptide “cocktails” is functionally important, but with plasticity over long evolutionary time scales.

## Introduction

The nervous systems of animals utilize a wide variety of chemicals for neuronal communication. These include amino acids (*e.g*. glutamate), biogenic amines (*e.g*. serotonin), and neuropeptides (*e.g*. vasopressin) amongst others. Neuropeptides are by far the most-diverse and they control many physiological/behavioural processes, including feeding, reproduction and locomotion [1–3]. Recent advances in genome/transcriptome sequencing are enabling identification of neuropeptide precursor proteins in species from a growing variety of animal taxa, providing new insights into the evolution of neuropeptide signaling [4–8]. The echinoderms are notable in this regard because as deuterostomian invertebrates they occupy an “intermediate” phylogenetic position with respect to the vertebrates and intensely studied protostomian invertebrates such as insects (*e.g. Drosophila melanogaster*) and nematodes (*e.g. Caenorhabditis elegans*). Accordingly, characterisation of neuropeptide signaling systems in echinoderms has recently provided key “missing links” for determination of neuropeptide relationships and reconstruction of neuropeptide evolution [8–10].

The phylum Echinodermata comprises five extant classes: Echinoidea (sea urchins and sand dollars), Holothuroidea (sea cucumbers), Asteroidea (starfish), Ophiuroidea (brittle stars and basket stars) and Crinoidea (sea lilies and feather stars). Recent molecular phylogenetic studies support the hypothesis that Echinoidea and Holothuroidea are sister groups (Echinozoa) and Asteroidea and Ophiuroidea are sister groups (Asterozoa), with the Crinoidea basal to the Echinozoa + Asterozoa clade (Eleutherozoa) [11, 12]. Echinoderms are marine organisms that have several unique features including pentaradial symmetry as adults, a remarkable ability to autotomise and regenerate body parts, and neurally-controlled mutable collagenous tissue [13, 14]. Previous transcriptomic analyses have identified neuropeptide precursor complements in *Strongylocentrotus purpuratus* (purple sea urchin), *Apostichopus japonicus* (Japanese sea cucumber) and *Asterias rubens* (common European starfish) [8, 15, 16]. Furthermore, the identification of neuropeptides in these species has facilitated investigation of the evolution and physiological roles of various neuropeptide signaling systems [8–10, 17–21].

The recent progress in transcriptomic/genomic characterization of echinoderm neuropeptide systems has hitherto not been extended to ophiuroids or crinoids. The Ophiuroidea constitutes the largest class among extant echinoderms [22] with a long evolutionary history that extends back to the early Ordovician (around 480 million years ago) [23], whilst extant families date from the mid-Permian (~ 270 million years ago) [12]. Available molecular data for ophiuroids has increased significantly in recent years with the emergence of numerous transcriptomic studies [20, 24–29]. Here, we utilize transcriptome sequence data from three brittle star species, *Ophionotus victoriae, Amphiura filiformis* and *Ophiopsila aranea* to perform the first comprehensive identification of neuropeptide precursors in ophiuroids. We identify representatives of over thirty neuropeptide families including homologs of endothelin/CCHamide, eclosion hormone (EH), neuropeptide-F/Y (NPF/NPY) and nucleobinin (NUCB)/nesfatin, which are the first to be discovered in a deuterostome/echinoderm.

Transcriptomes have also been employed to investigate the phylogenetic relationships of the ophiuroids, utilising data from fifty-two species [12]. In this the most comprehensive molecular analysis of ophiuroid phylogeny to date, previous morphology-based classification schemes [30] were rejected in favour of a new phylogeny comprising three primary ophiuroid clades [12, 31, 32]. This landmark study and the associated large dataset has provided a unique opportunity to investigate the conservation and diversification of neuropeptide precursor structure over a period of ~270 million years of ophiuroid evolution. Our analysis reveals that the majority of ophiuroid neuropeptide precursors contain a single copy of a neuropeptide, but several precursors comprise multiple copies of identical or non-identical, but structurally-related, neuropeptides. Interestingly, the number of neuropeptide copies in the majority of precursors is constant across all the ophiuroid species examined, but examples of clade-specific losses/gains of neuropeptides are also observed. This remarkable conservation in neuropeptide copy number across ~270 million years of ophiuroid evolution indicates that the composition of neuropeptide “cocktails” is functionally important, but with plasticity over long evolutionary time scales.

## Results and discussion

Here, we have utilized transcriptome sequence data for the first comprehensive identification of neuropeptide precursors in ophiuroids (**Figure 1).** Representatives of over thirty bilaterian neuropeptide precursor families were identified. Identification of ophiuroid representatives of these neuropeptide precursor types has in some cases provided new insights into neuropeptide precursor structure and evolution, as discussed in more detail below. First, however, we will highlight representatives of bilaterian neuropeptide precursor families that have been identified here for the first time in an echinoderm species.

**Figure 1.**
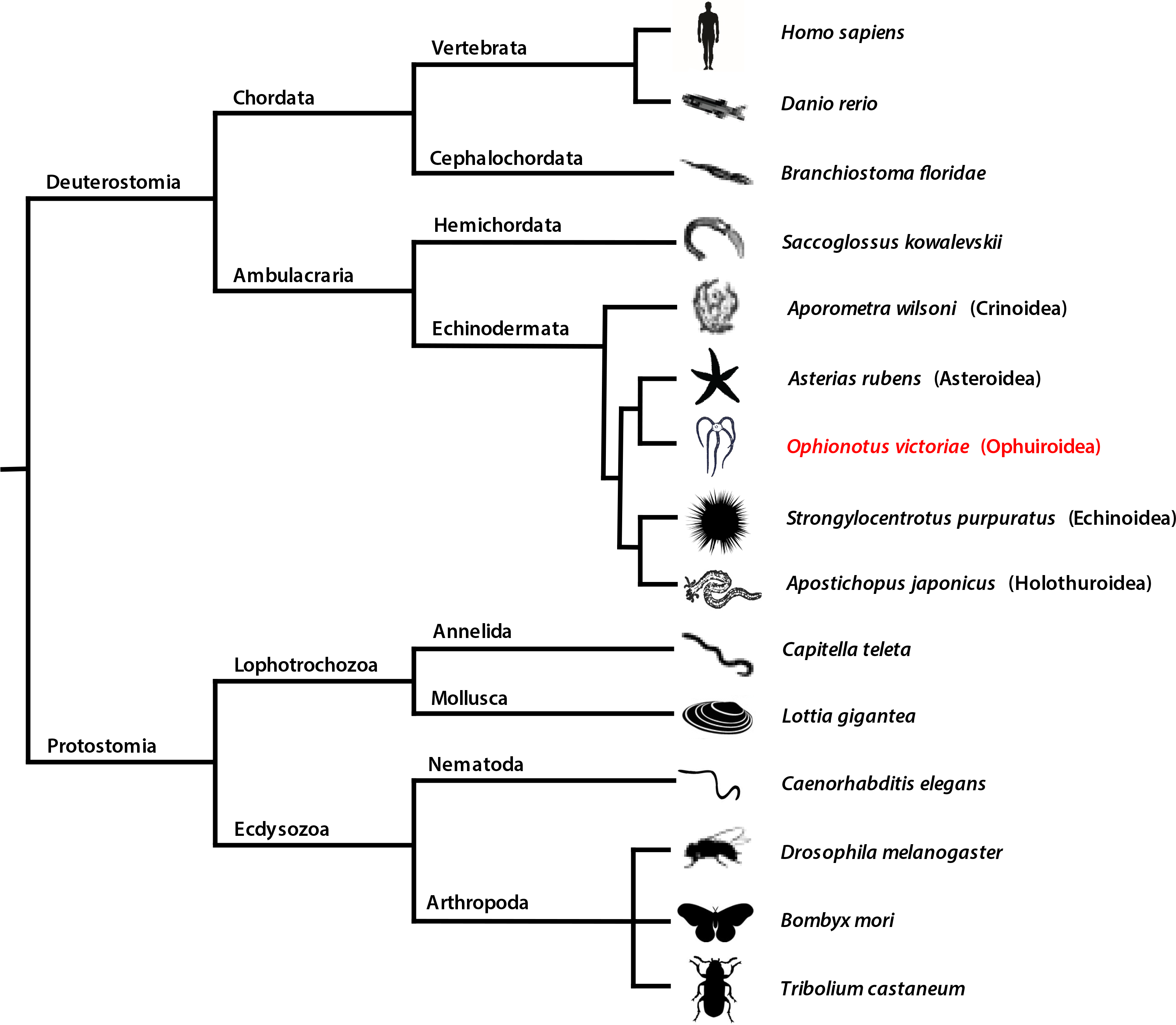
Bilaterian animal phylogeny. The diagram shows i). the phylogenetic position of the phylum Echinodermata in the ambulacrarian clade of the deuterostomes and ii) relationships between the five extant classes of echinoderms, which include the focal class for this study–the Ophiuroidea (e.g. *Ophionotus victoria*e).

### Discovery of the first echinoderm representatives of bilaterian neuropeptide families

Comprehensive analysis of transcriptome sequence data from three ophiuroid species, *O. victoriae, A. filiformis and O. aranea*, has enabled the discovery of the first echinoderm representatives of four bilaterian neuropeptide families. Specifically, we have discovered the first deuterostomian homologs of eclosion hormone (**Figure 2**), the first ambulacrarian homolog of CCHamide/endothelin-type peptides (**Figure 3A**), and first echinoderm homologs of neuropeptide-Y/neuropeptide-F (**Figure 3B**) and NUCB/nesfatin (**Figure S1**), as discussed in detail below.

**Figure 2.**
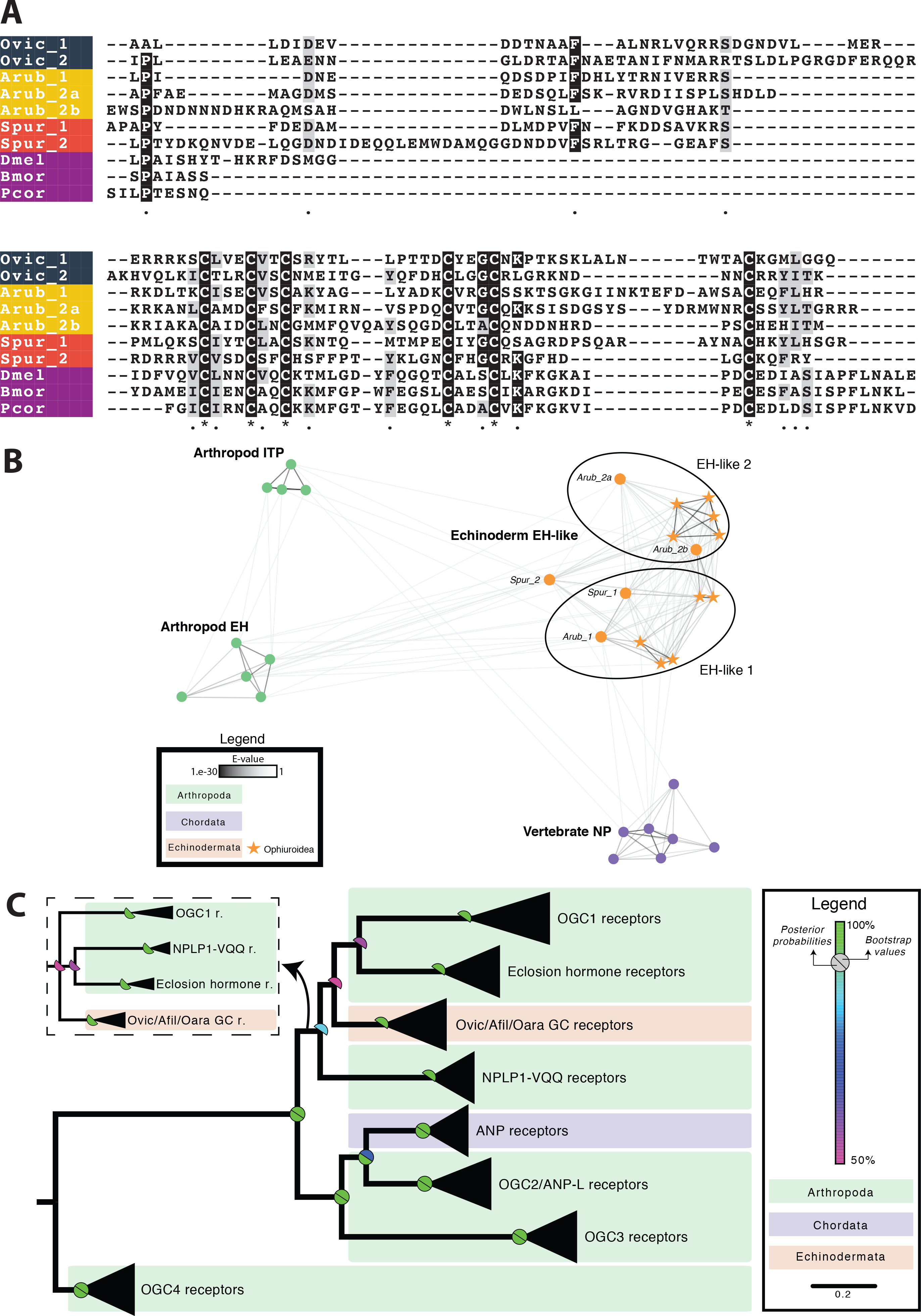
Eclosion hormone (EH)-type peptides and receptors in echinoderms A) Partial multiple sequence alignment of eclosion hormone-type precursor sequences, excluding the N-terminal signal peptide; B) Cluster analysis of arthropod EH precursors, echinoderm EH-like precursors, arthropod ion transport peptides (ITPs) and vertebrate atrial natriuretic peptides shows that echinoderm EH-like precursors are more closely related to arthropod EH than ITP C) Maximum likelihood and Bayesian phylogenetic analyses of membrane guanylate cyclase receptors shows that EH-like receptors are found in echinoderms but are absent in vertebrates as seen for the EH-like precursors. OGC1, 2, 3 and 4 are orphan guanylate cyclase receptors found in arthropods [42]. Echinoderm EH-like receptors are clustered with arthropod EH receptors, neuropeptide-like peptide 1-VQQ receptors (NPLP1-VQQ) and OGC1 receptors. The inset shows the alternate topology obtained following Bayesian analysis. Species names: *Ophionotus victoriae* (Ovic), *Asterias rubens* (Arub), *Strongylocentrotus purpuratus* (Spur), *Drosophila melanogaster* (Dmel), *Bombyx mori* (Bmor) and *Pediculus humanus corporis* (Pcor).

**Figure 3.**
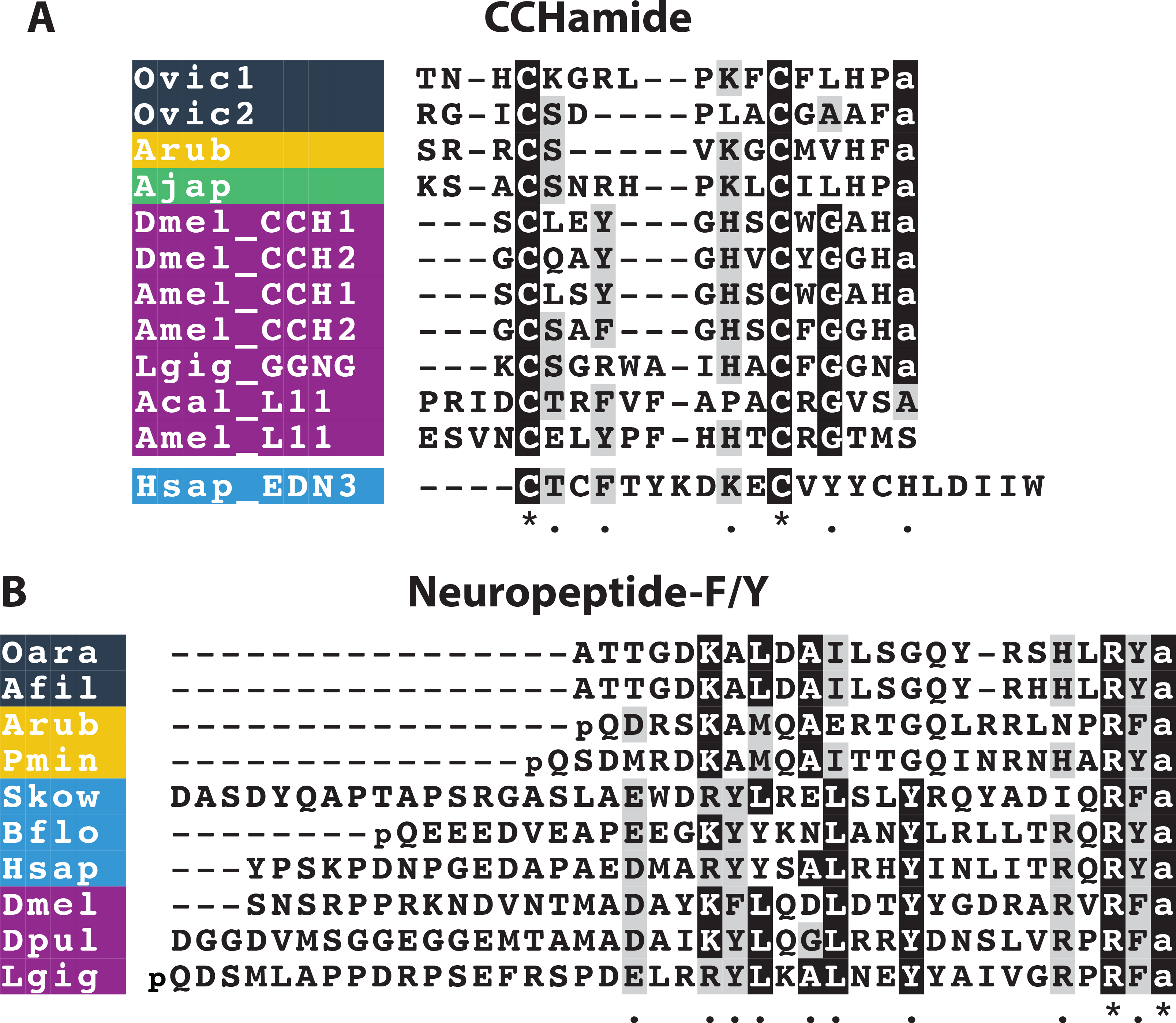
Multiple sequence alignments of A) CCHamide-type and B) Neuropeptide-F/Y-type peptides. Species names: *Ophionotus victoriae* (Ovic), *Asterias rubens* (Arub), *Apostichopus japonicus* (Ajap), *Drosophila melanogaster* (Dmel), *Apis mellifera* (Amel), *Lottia gigantea* (Lgig), *Aplysia californica* (Acal), *Homo sapiens* (Hsap), *Ophiopsila aranea* (Oara), *Amphiura filiformis* (Afil), *Patiria miniata* (Pmin), *Saccoglossus kowalevskii* (Skow), *Branchiostoma floridae* (Bflo) and *Daphnia pulex* (Dpul).

#### Eclosion hormone

Eclosion hormone (EH) was first isolated and sequenced in the insects *Manduca sexta* (tobacco hornworm) and *Bombyx mori* (silk moth) and shown to alter the timing of adult emergence [33, 34]. EH is one of the main peptide/protein hormones involved in control of ecdysis (*i.e*. shedding of the cuticle) behavior in arthropods [35, 36]. It binds to and activates a receptor guanylyl cyclase that is expressed in epitracheal Inka cells and causes the secondary release of ecdysis-triggering hormone (ETH) that is also expressed in Inka cells [37, 38]. In *Drosophila*, EH is important for ecdysis but whether this hormone is essential for ecdysis is not yet clear [39, 40]. EH null mutant flies show defects in ecdysis and are unable to reach adulthood yet some flies in which EH-producing neurons have been genetically ablated (a more extreme manipulation) are able to survive till adulthood. Arthropod EHs have six conserved cysteine residues that form three disulfide bridges [37]. EHs have not been discovered previously outside of arthropods. Interestingly, four EH-like precursors were identified in *A. filiformis* and *O. aranea* and two in *O. victoriae* (**Figure S2–S4**, GenBank; MF155236; MF155237). The ophiuroid EH-like precursors are orthologous to neuropeptide precursors previously identified in the sea-urchin *S. purpuratus* (Spnp11 and Spnp15, which we now rename as Spur EH1 and Spur EH2, respectively) [16] and the starfish *A. rubens* (Arnp11, Arnp15 and Arnp15b renamed as Arub EH1, Arub EH2a and Arub EH2b, respectively) [8]. The positions of cysteine residues are conserved across all echinoderm and insect EHs, but aside from this there is little sequence conservation (**Figure 2A**). The echinoderm EH-like precursor sequences were also analysed using a sequence-similarity-based clustering approach based on BLASTp e-values using CLANS software [41]. The analysis shows that echinoderm EH-like precursors (i) cluster in two compact subgroups (echinoderm EH-like precursor 1 and EH-like precursor 2 and (ii) have strong positive BLAST results with arthropod EHs and, to a lesser extent, with arthropod ion transport peptide (ITP) and vertebrate atrial natriuretic peptide (ANP) (**Figure 2B**). ITP precursors also possess six cysteine residues; however, the position of these residues is not conserved with cysteine residues found in echinoderm EH-like precursors (not shown).

To obtain further evidence for the presence of an EH-like signaling system in echinoderms, we performed a phylogenetic analysis of EH-type receptors. Insect EHs mediate their effects by binding to membrane guanylyl cyclase receptors [38]. EH receptors are closely related to vertebrate ANP receptors and various orphan receptors [42]. Specific BLAST searches enabled identification of transcripts in *O. victoriae*, *A. filiformis* and *O. aranea* that encode proteins similar to arthropod EH receptors. Maximum likelihood and Bayesian phylogenetic analyses confirmed that these sequences group with the receptor cluster containing EH receptors (**Figure 2C**). The discovery of the first deuterostomian EHs suggests an ancient bilaterian origin of EHs and indicates that these hormones may have other functions in invertebrates aside from their role in ecdysis.

#### CCHamide

CCHamides are neuropeptides that were discovered relatively recently in the silkworm *Bombyx mori* [43]. Later, it was found that insects have two CCHamide genes, CCHamide-1 and CCHamide-2, each encoding a single copy of the mature peptide [44]. These peptides are referred to as CCHamides because they contain two cysteine residues and a characteristic histidine-amide C-terminal motif. There are two CCHamide receptors in insects: CCHamide-1 specifically activates one receptor and CCHamide-2 specifically activates the second receptor [44, 45]. CCHamide-1 has a physiological a role in starvation-induced olfactory modifications [46] whereas as CCHamide-2 regulates feeding, growth and developmental timing in flies [45, 47]. Recent studies examining the evolution of neuropeptides in the Bilateria have shown that protostomian CCHamides are related to elevenin (another protostomian neuropeptide originally discovered from the mollusc *Aplysia californica* L11 neuron), lophotrochozoan GGNG peptides, endothelins and gastrin-releasing peptides (GRPs) [6, 7, 48, 49]. The latter two are neuropeptide types that have not been found outside chordates. Furthermore, the degree of sequence/structural conservation varies across these different peptide families. Hence, CCHamides are amidated and have a disulphide bridge, elevenins and endothelins have a disulphide bridge but are non-amidated and GRPs are amidated but lack the disulphide bridge. Furthermore, CCHamide-1 is located immediately after the signal peptide whereas there is a dibasic cleavage site separating the signal peptide and CCHamide-2 [44].

Here we have identified two neuropeptide precursors in brittle stars whose sequence and precursor structure resembles those of lophotrochozoan GGNG peptides and insect CCHamide-1 (**Figure 3A**). The CCHamide-like precursor 1 (GenBank; MF155229) identified in *O. victoriae* is orthologous to an uncharacterized neuropeptide precursor (Arnp25) identified previously in the starfish *A. rubens* [8], whereas the CCHamide-like precursor 2 (GenBank; MF155230) was only found in brittle stars. Both CCHamide-like precursors in *O. victoriae* comprise a single copy of a putative cyclic amidated peptide that is flanked by a signal peptide at the N-terminus and a dibasic cleavage site at the C-terminus. Interestingly, both of these peptides lack a penultimate histidine residue, just like the lophotrochozoan GGNG peptides (**Figure 3A**) [48, 49].

#### Neuropeptide-Y/Neuropeptide-F

Neuropeptide-Y (NPY) was first isolated and sequenced from the porcine hypothalamus in 1982 [50, 51]. Although the NPY/NPF family of peptides are pleiotropic in nature [52], they are mainly known for their roles in regulation of feeding and stress [3, 53, 54]. The discovery of Neuropeptide-F (NPF) in the tapeworm *Monieza expansa* in 1991 demonstrated for the first time the occurrence of NPY homologs in invertebrates [55]. Here, we have identified the first echinoderm representatives of the NPY/NPF family in brittle stars and starfish (**Figure 3B, Figure S12**). The brittle star precursors contain a peptide with a C-terminal RYamide, in common with NPY in vertebrates and an ortholog in the starfish *Patiria miniata*. In contrast, an ortholog in the starfish *A. rubens* has a C-terminal RFamide, a feature that it shares with NPY/NPF-type peptides in the hemichordate *S. kowalevskii* and in protostomes. Thus, our findings have revealed that NPY/NPF-type peptides with a C-terminal Yamide motif are not restricted to vertebrates, as has been shown previously in some insects [52]. Echinoderm NPY/NPF-type peptides are located immediately after the signal peptide in the precursor proteins, as is the case in other bilaterian species. Surprisingly, we did not find NPY/NPF-type precursors in the sea urchin *S. purpuratus* or the sea cucumber *A. japonicus*. However, we suspect that this may reflect sequence divergence rather than gene loss because a gene encoding a NPY/NPF-type receptor can be found in the *S. purpuratus* genome [56].

#### NUCB

Nucleobindins (NUCB1 and NUCB2) are multidomain Ca^2+^ and DNA binding proteins. NUCB1 was first discovered in 1992 and thought to play a role in apoptosis and autoimmunity [57]. Interestingly, the NUCB1 precursor has both a signal peptide and a leucine zipper structure suggesting that it can bind DNA and act as an endocrine factor [58]. NUCB2 is a homolog of NUCB1 and was named based on high sequence similarity between the two precursors [59]. In 2006, an 82 amino acid peptide located in the N-terminal region of NUCB2 was reported. This peptide, Nesfatin-1 (Nucleobindin-2-Encoded Satiety and FAT-Influencing proteiN-1), was discovered as a satiety inducing factor in the rat hypothalamus [60]. Its role in inhibiting food intake in vertebrates is now well-established [59, 61]. Moreover, this pleiotropic peptide also modulates other processes including glucose and lipid metabolism, and cardiovascular and reproductive functions. Recently, nesfatin-1-like peptide derived from NUCB1 was shown to be anorexigenic in goldfish [62]. Surprisingly, the presence of NUCBs in invertebrates other than *Drosophila* has not been reported until now [63]. Here, we show that NUCB-type precursors are present in echinoderms (**Figure S1A**). Phylogenetic analysis of NUCB precursors reveals that a single copy of the NUCB precursor is found in invertebrate species and gene duplication in the vertebrate lineage gave rise to NUCB1 and NUCB2 (**Figure S1B**). In chordates, the NUCB precursors are predicted to generate three peptides (Nesfatin-1, 2 and 3); however, no biological role has been attributed specifically to nesfatin-2 and nesfatin-3. Interestingly, the prohormone convertase cleavage sites expected to generate Nesfatin-1, 2 and 3 are conserved between echinoderm and chordate NUCBs. Moreover, the *O. victoriae* precursor (Genbank; MF155235) has an additional predicted cleavage site within the Nesfatin-1 containing region, which is not present in other species (except for *Drosophila melanogaster*). However, it remains to be determined whether or not this cleavage site in the *O. victoriae* precursor is functional.

### First comprehensive identification of neuropeptide precursors in ophiuroids

We have identified neuropeptide precursors belonging to 32 families, which represents the first comprehensive analysis of neuropeptide precursors in ophiuroids (**Figure 4; Figure S2–S4**). Several of these neuropeptide families have been identified previously in echinoderms and include homologs of AN peptides, bursicon (α and β) (GenBank; MF155260; MF155227), calcitonin (GenBank; MF155228), cholecystokinin (CCK) (GenBank; MF155231; MF155232) [15], corazonin (GenBank; MF155233) [10], corticotropin-releasing hormone (CRH) (GenBank; MF155234; MF155235, MF155261, MF155262), glycoprotein hormones (α2 and β5) (GenBank; MF155238; MF155239; MF155240) [64], gonadotropin-releasing hormone (GnRH) (GenBank; MF155263) [10], insulin-like peptide (ILP) (GenBank; MF155264) [64], kisspeptin (KP) (GenBank; MF155241) [8], luqin (GenBank; MF155242) [7], melanin-concentrating hormone (MCH) (GenBank; MF155243) [8], NG peptides (neuropeptide-S) (GenBank; MF155244) [9, 65], orexin (GenBank; MF155245; MF155246) [6, 8], pedal peptides (GenBank; MF155247; MF155266; MF155267) [16], pigment-dispersing factor (PDF) (GenBank; MF155248) [8], relaxin-like peptide (GenBank; MF155249) [66], SALMFamides (L-type and F-type) (GenBank; MF155250; MF155268) [19, 20, 67], somatostatin (GenBank; MF155252; MF155253) [8], tachykinin (GenBank; MF155254) [8], thyrotropin-releasing hormone (TRH) (GenBank; MF155255; MF155256) [16] and vasopressin/oxytocin (GenBank; MF155257) [64, 65] (**Figures 5–7 and S5-S10**). With the exception of MCH (which may be unique to deuterostomes) [6, 8], AN peptides and SALMFamides (which thus far have only been identified in echinoderms), the origins of all of the neuropeptide precursors identified here in ophiuroids predate the divergence of protostomes and deuterostomes [6, 7]. Of the three species examined here, the neuropeptide precursor complement of *O. victoriae* was the most complete (**Figure 4**) and therefore this species is used as a representative ophiuroid for sequence alignments, except in a few cases where a neuropeptide precursor was not found in *O. victoriae*. Below we highlight several interesting and/or unusual features of ophiuroid neuropeptides and neuropeptide precursors.

**Figure 4.**
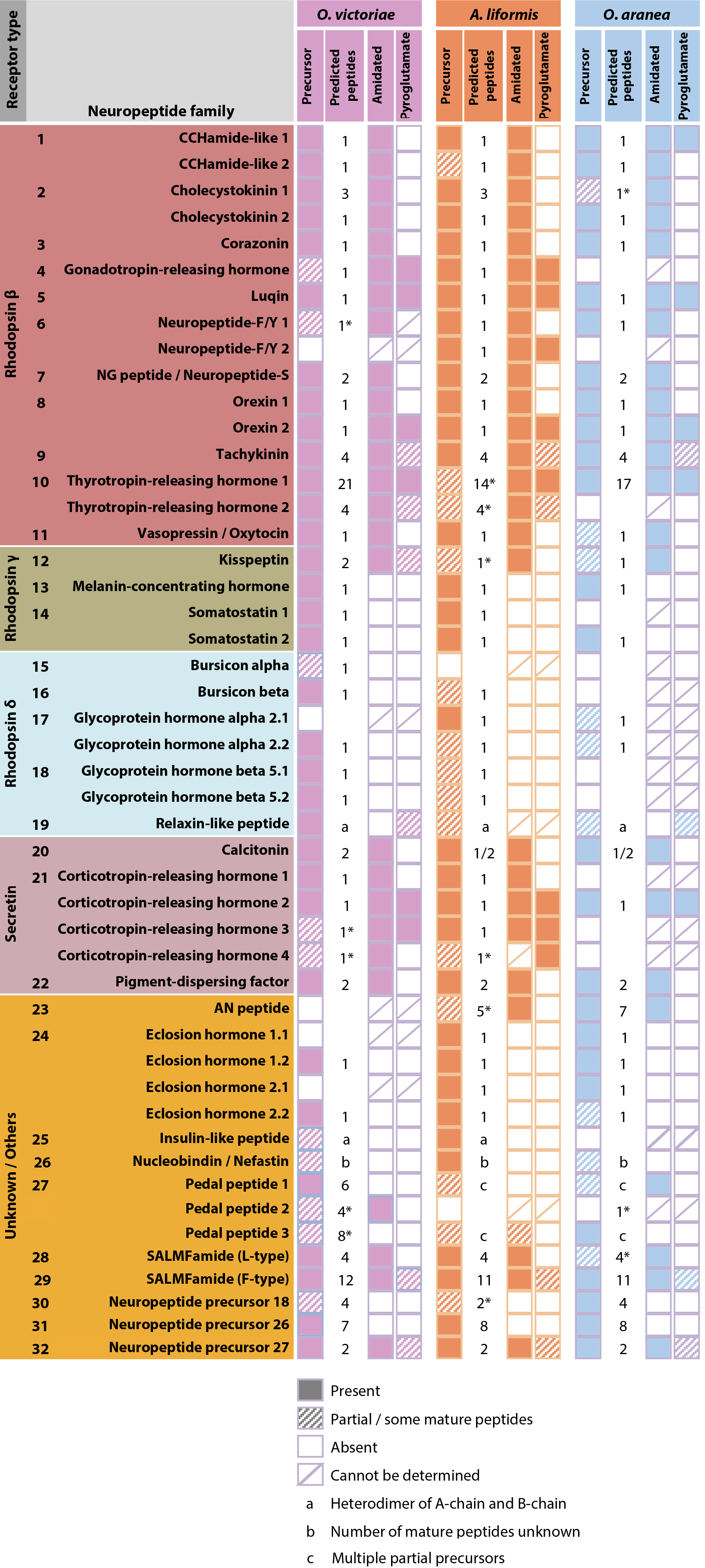
Summary of neuropeptide precursors identified in *Ophionotus victoriae, Amphiura filiformis* and *Ophiopsila aranea*. Neuropeptide precursors are classified based on the type of G-protein coupled receptor (GPCR) their constituent peptides are predicted to activate (see Mirabeau and Joly, 2013). Some peptides bind to receptors other than GPCRs and these are grouped with peptides where the receptor is unknown. Ophiuroids have neuropeptide precursors from up to 32 families. The number of putative mature peptides derived from each precursor has been indicated along with the presence of amidation and pyroglutamation.

**Figure 5.**
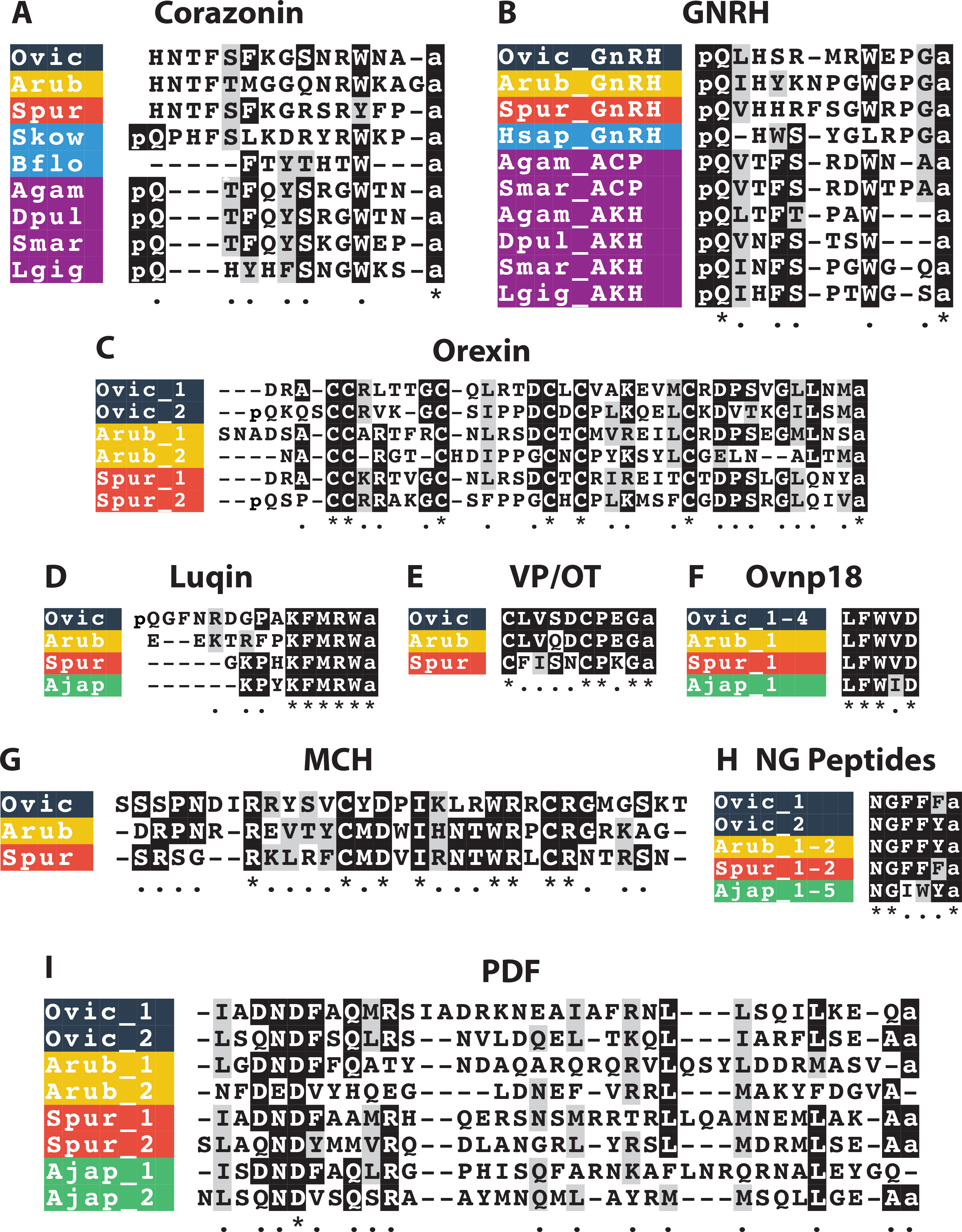
Multiple sequence alignments of mature peptides belonging to selected neuropeptide families. A) corazonin alignment; B) gonadotropin-releasing hormone (GnRH) alignment; C) orexin alignment; D) luqin alignment; E) vasopressin/oxytocin (VP/OT) alignment; F) Ovnp18 alignment; G) melanin-concentrating hormone (MCH) alignment; H) NP peptide alignment; I) pigment dispersing factor (PDF) alignment (see Figure S10 for a multiple sequence alignment of PDF-type precursors). Species names: *Ophionotus victoriae* (Ovic), *Asterias rubens* (Arub), *Strongylocentrotus purpuratus* (Spur), *Apostichopus japonicus* (Ajap), *Saccoglossus kowalevskii* (Skow), *Branchiostoma floridae* (Bflo), *Anopheles gambiae* (Agam), *Daphnia pulex* (Dpul), *Strigamia maritima* (Smar), *Lottia gigantea* (Lgig) and *Homo sapiens* (Hsap).

**Figure 6.**
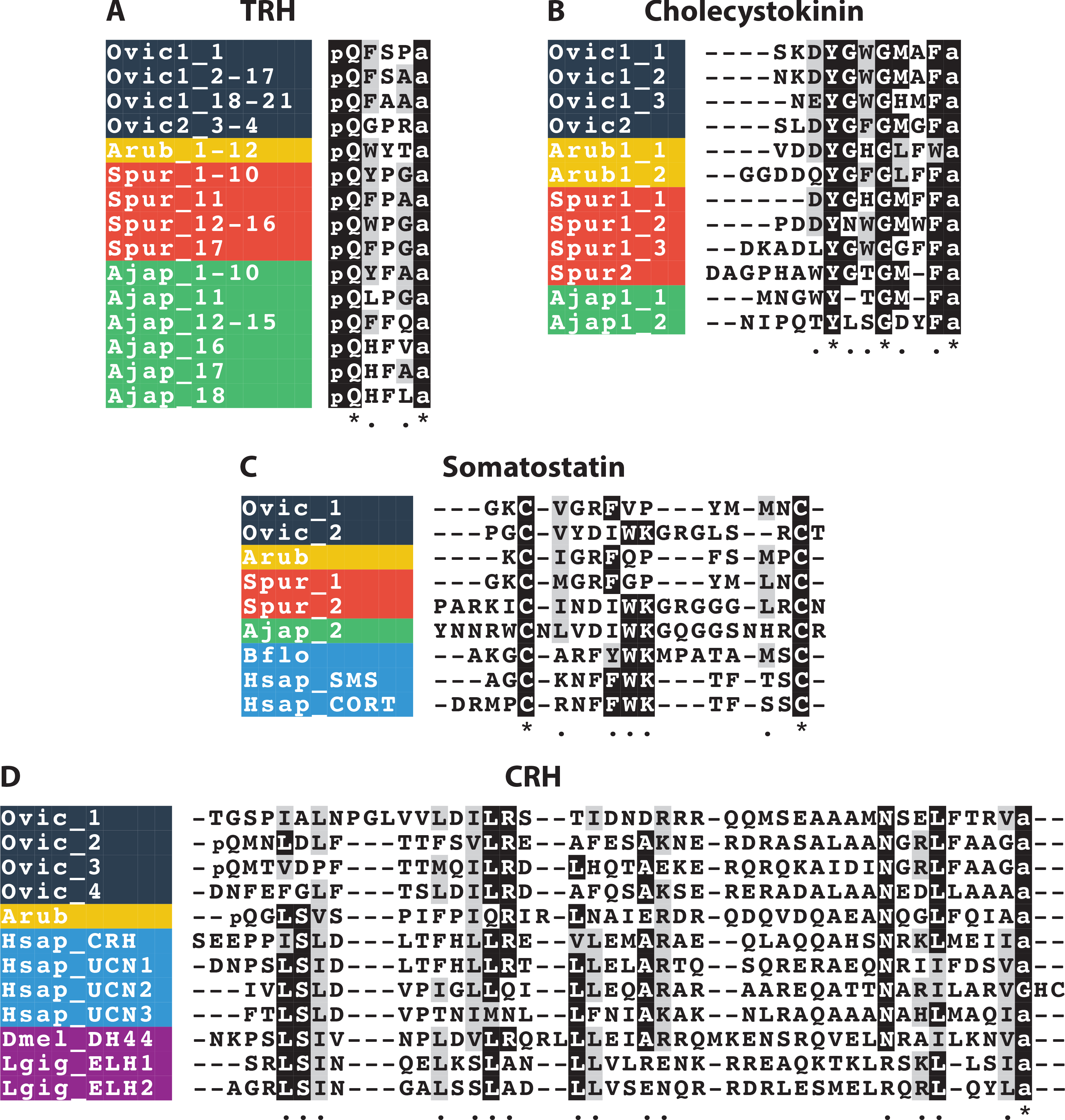
Alignments of neuropeptides derived from precursors that exist in multiple forms in ophiuroids. A) thyrotropin-releasing hormone (TRH) alignment; B) cholecystokinin alignment; C) somatostatin alignment; D) corticotropin-releasing hormone (CRH) alignment. Species names: *Ophionotus victoriae* (Ovic), *Asterias rubens* (Arub), *Strongylocentrotus purpuratus* (Spur), *Apostichopus japonicus* (Ajap), *Branchiostoma floridae* (Bflo), *Homo sapiens* (Hsap), *Drosophila melanogaster* (Dmel) and *Lottia gigantea* (Lgig).

**Figure 7.**
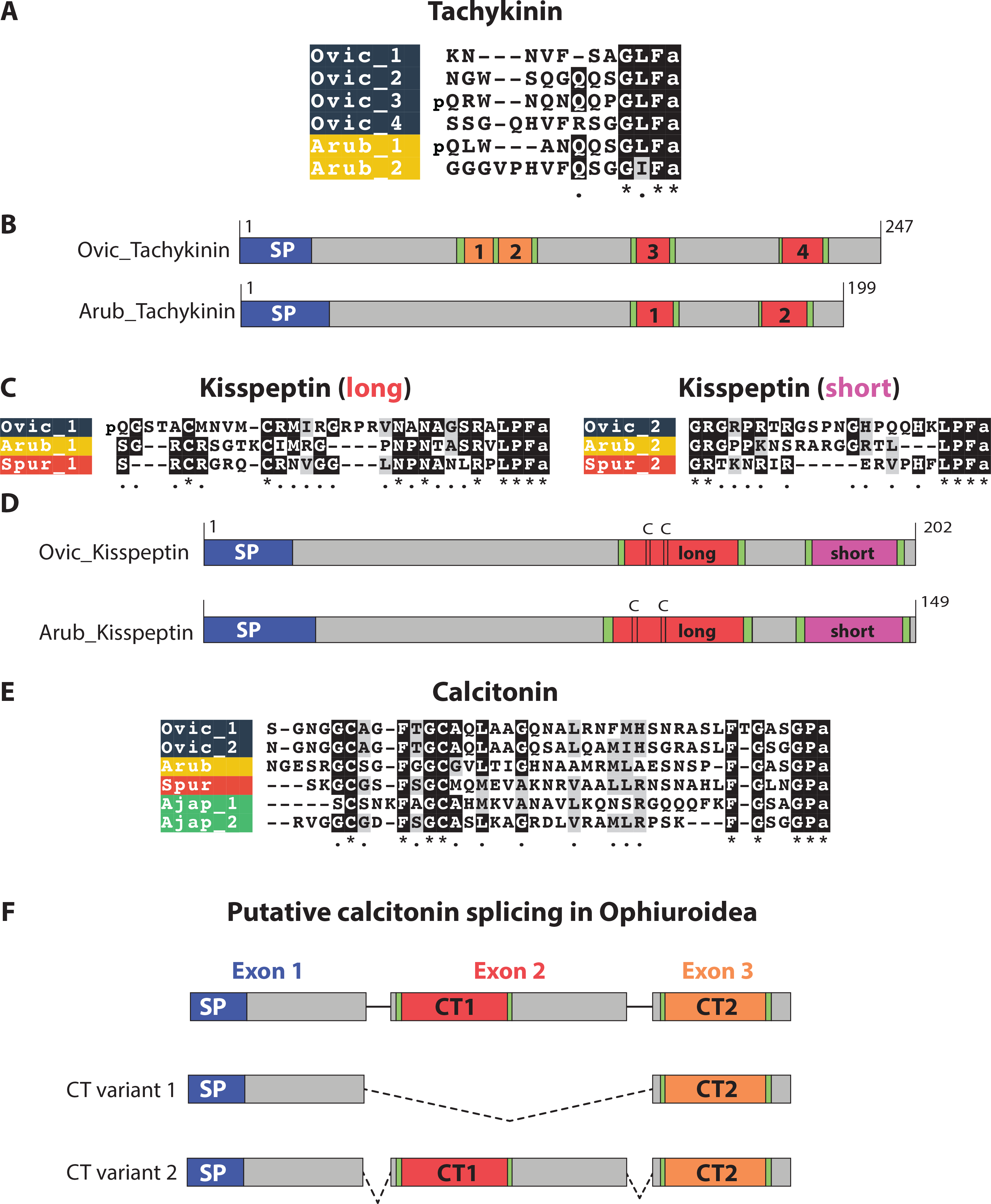
Comparative analysis of ophiuroid tachykinin, kisspeptin and calcitonin-type precursors and neuropeptides. A) Alignment of tachykinin-type peptides in *O. victoriae* (Ophiuroidea) and *A. rubens* (Asteroidea); B) Schematic diagrams of the *O. victoriae* and *A. rubens* tachykinin precursors showing the location of the signal peptide (SP) and predicted neuropeptides (labelled 1 to 4); C) Alignments of the long and short forms of kisspeptin-type neuropeptides in *O. victoriae, A. rubens* and *S. purpuratus* (Echinoidea) D) Schematic diagrams of the *O. victoriae* and *A. rubens* kisspeptin precursors showing the locations of the SP, short and long orthocopies and cysteine (C) residues; E) Alignment of calcitonin-type peptides from *O. victoriae, A. rubens, S. purpuratus* and *A. japonicus* (Holothuroidea); F) Predicted alternative splicing of the calcitonin gene in ophiuroids, with the location of the SP and neuropeptides (CT1 and CT2) labelled. Species names: *Ophionotus victoriae* (Ovic), *Asterias rubens* (Arub), *Strongylocentrotus purpuratus* (Spur) and *Apostichopus japonicus* (Ajap).

### Neuropeptide precursors that occur in multiple forms in O. victoriae

#### Thyrotropin-releasing hormone (TRH)-type precursors

TRH (also known as thyrotropin-releasing factor or thyroliberin) was first isolated and sequenced in the 1960s [68–70]. In mammals, TRH is produced in the hypothalamus and stimulates the release of thyroid-stimulating hormone (TSH) and prolactin from the anterior pituitary [71, 72]. The recent discovery of a TRH receptor in the annelid *Platynereis dumerilii* indicates that the evolutionary origin of this neuropeptide signaling system predates the divergence of protostomes and deuterostomes [73].

The human TRH precursor contains six copies of the tripeptide pQHPamide [74]. Precursor proteins comprising multiple copies of TRH-like peptides have been identified previously in the sea urchin *S. purpuratus*, the sea cucumber *A. japonicus* and the starfish *A. rubens* [8, 15, 16], with a single TRH-type precursor found in each of these species. Interestingly, here we identified two TRH-type precursors (OvTRHP1 and OvTRHP2) in *O. victoriae* (**Figure S2 and 6A**). OvTRHP1 comprises 21 copies of putative TRH-like tetrapeptides with the motif pQXXXamide (where X is variable). OvTRHP2, on the other hand, comprises two copies of the putative tetrapeptide pQGPRamide and two longer peptides that also have a C-terminal GPRamide motif but lack the N-terminal pyroglutamate.

#### Cholecystokinin (CCK)-type precursors

A CCK-type peptide (formerly pancreozymin) was first sequenced in the 1960s [75]. CCK-type peptides play numerous roles in feeding and digestion related physiology. CCK mediates satiety, stimulates the release of digestive enzymes and gall bladder contractions [76–78]. CCK-type peptides are involved in mechanisms of learning and memory, and analgesia [79]. A neuropeptide precursor comprising two CCK-like peptides was recently identified in the starfish *A. rubens* [8]. Here we have identified two CCK-type precursors in *O. victoriae* (OvCCKP1 and OvCCKP2) and orthologs of both of these precursors were also identified in the sea urchin *S. purpuratus* (**Figure S2**) [16]. The CCK-type precursor 1 comprises three CCK-like peptides in both *O. victoriae* and *S. purpuratus* and this precursor is similar to the *A. rubens* CCK-type precursor, which comprises two CCK-like peptides. In contrast, the CCK-type precursor 2 comprises a single CCK-like peptide in both *O. victoriae* and *S. purpuratus*. Interestingly, the sequence of the *S. purpuratus* CCK-type precursor 2 was reported previously as part of a genome-wide search for neuropeptides [80], but the authors of this study did not identify it as a CCK-type precursor. However, based on the presence of a conserved tyrosine residue and a C-terminal F-amide motif in the predicted neuropeptide derived from this protein, it is evident that it belongs to the family of CCK-type precursors (**Figure 6B**). A search of a preliminary genome assembly of the starfish *Patiria miniata* (http://www.echinobase.org) [81] did not reveal a gene encoding a CCK-type precursor 2. Therefore, it appears that this neuropeptide precursor type may have been lost in the Asteroidea; nevertheless, further analysis of a wider range of starfish species will be required to draw definitive conclusions. With a broader evolutionary perspective, CCK-type peptides in deuterostomes are orthologs of sulfakinin (SK)-type neuropeptides found in insects [6, 7]. Interestingly, insects have a single SK precursor, which comprises two neuropeptides, SK-1 and SK-2 [82], and this may reflect the ancestral condition in the common ancestor of protostomes and deuterostomes. Thus, the occurrence of two CCK-type peptides on a single precursor in *A. rubens* and insects may be an ancestral characteristic and the occurrence of two CCK-type precursors that comprise one and three CCK-type peptides appears to be a derived characteristic.

#### Somatostatin-type precursors

Somatostatin was first isolated and sequenced from sheep hypothalamus in 1973 [83]. This peptide inhibits the release of pituitary hormones such as growth hormone, prolactin and thyroid-stimulating hormone [84]. Moreover, it also inhibits the release of gastrointestinal (cholecystokinin and gastrin amongst others) and pancreatic (insulin and glucagon) hormones [85–87]. Aside from its effects on release of hormones, somatostatin also has central actions that influence motor activity [85]. Here, we have identified two somatostatin-type precursors (OvSSP-1 and OvSSP-2) in *O. victoriae*. (**Figure S2 and 6C**). Homologs of both of these precursors are present in the sea urchin *S. purpuratus* (**Figure S2 and 6C**), one of which was previously referred to as Spnp16 [16]. By comparison, only a single somatostatin-type precursor has been found in the starfish *A. rubens*, which is an ortholog of OvSSP-1 [8]. All somatostatin-type precursors comprise a single copy of the bioactive neuropeptide, which is located in the C-terminal region of the precursor [88, 89]. Interestingly, the type-1 somatostatins in echinoderms have a phenylalanine residue located in the middle part of the peptide and this conserved feature is found in human somatostatin. Conversely, type-2 somatostatins in echinoderms lack the phenylalanine residue but have a neighbouring tryptophan-lysine (WK) motif that is also conserved in human and *B. floridae* somatostatins (**Figure 6C**). The deuterostomian somatostatins are orthologous to the allatostatin-C neuropeptide family in arthropods [7]. This family of peptides comprises three precursor-types: allatostatin-C, allatostatin-CC and the recently discovered allatostatin-CCC [89, 90]. Both allatostatin-C and allatostatin-CC are non-amidated, like somatostatins; however, allatostatin-CCC has a C-terminal amide. Hence, non-amidated peptides may be representative of the ancestral condition in the common ancestor of protostomes and deuterostomes, with the amidated allatostatin-CCC probably having evolved only within the arthropod lineage [90]. It remains to be determined whether or not the duplication of somatostatin-type precursors in echinoderms and the duplication of allatostatin C (to give rise to allatostatin-CC) represent independent duplications. Further insights into this issue may be obtained if the receptors for somatostatin-type peptides in echinoderms are deorphanised.

#### CRH-type Corticotropin-releasing hormone

(CRH)-type precursors peptides are a family of related neuropeptides that include CRH, urocortins and urotensin-I in chordates, egg-laying hormone (ELH) in lophotrochozoans and diuretic hormone 44 (DH_44_) in arthropods [6, 7]. Arthropods usually have a single DH_44_ precursor, which comprises a single copy of the mature peptide. In some insects, such as *Tribolium castaneum* and *Bombyx mori*, alternative splicing of DH_44_ transcripts results in multiple mature peptide isoforms of varying lengths [43, 91]. The situation in lophotrochozoans is more complex, with several species having multiple precursors and some of these precursors comprising multiple ELH mature peptides [4, 92]. A single CRH-type precursor was found previously in the starfish *A. rubens*, whereas here we have identified four CRH-type precursors in *O. victoriae* (**Figure S2 and 6D**). Thus, expanded families of CRH-type peptides and receptors appear to have evolved independently in multiple animal lineages, including chordates and ophiuroid echinoderms [93, 94].

### Diversity in neuropeptide precursor structure: new insights from ophiuroids

#### Tachykinins

The mammalian neuropeptide substance P was the first tachykinin-type peptide to be isolated and sequenced [95–97]. Subsequently, tachykinin-type peptides were discovered in other animals including tunicates [98], insects [99, 100], annelids [101] and molluscs [102]. Tachykinin-type peptides regulate various physiological processes including muscle contractility [103], nociception [104] and stress responses [105] amongst others [106]. Analysis of genomic/transcriptomic sequence data from the sea urchin *S. purpuratus* and the sea cucumber *A. japonicus* did not identify candidate tachykinin-type precursors [6, 7, 15, 16]. However, recently a putative tachykinin-type precursor was discovered in the starfish *A. rubens* (ArTKP), indicating that this signaling system does occur in some echinoderms [8]. Here we have identified orthologs of ArTKP in *O. victoriae* and other ophiuroids (**Figure 4** and **7A**). Collectively, these findings indicate that this signaling system has been retained in the Asterozoa but lost in the Echinozoa. Comparison of the structure of the asterozoan tachykinin-type precursors reveals that the *A. rubens* precursor (ArTKP) comprises two putative mature peptides, whereas the *O. victoriae* precursor comprises four mature peptides (**Figure 7B**). It remains to be determined, however, which of these two conditions represents the ancestral state in the common ancestor of the Asterozoa. Further insights into this issue may be obtained if sequence data from a variety of starfish species are analysed.

#### Kisspeptins (KP)

Kisspeptin (formerly known as metastin) is encoded by the *KiSS1* gene in humans. *KiSS1* was originally discovered as a gene that may suppress the metastatic potential of malignant melanoma cells [107]. Subsequently, it was found to play a vital role in regulating the onset of puberty. Thus, in vertebrates kisspeptin binds to its receptor GPR54 to stimulate pituitary release of gonadotropin-releasing hormone (GnRH) [108]. The first KP-type precursors to be identified in non-chordates were discovered recently in ambulacrarians - the echinoderms *A. rubens* and *S. purpuratus* and the hemichordate *S. kowalevskii* [8]. Accordingly, here we have identified KP-type precursors in *O. victoriae* and other ophiuroids. All of the ambulacrarian precursor proteins comprise two KP-type peptides and the first putative neuropeptide in the echinoderm precursors has two cysteine residues at the N-terminus, which could form an N-terminal disulphide bridge similar that of calcitonin-type peptides (see below). In contrast, the second putative neuropeptide does not contain any cysteine residues and is typically shorter than the first peptide (**Figure 7C and D**). Interestingly, comparison of the sequences of the first (long) and second (short) KP-type peptides in echinoderms reveals that the long and short peptides share less sequence similarity with each other within a species than they do with respective peptides in other species (**Figure 7C**). This indicates that the duplication event that gave rise to the occurrence of the long and short peptides occurred before the divergence of the Asterozoa and Echinozoa. Interestingly, previous studies have revealed that there has been an expansion of KP-type receptors in ambulacraria (*S. purpuratus* and *S. kowalevskii*) and in the cephalochordate, *Branchiostoma floridae*, with 16 KP receptors present in the latter [6, 56]. Further studies are now needed to identify the proteins that act as receptors for the KP-type peptides identified here in ophiuroids and previously in other echinoderms [8].

#### Calcitonin

Calcitonin was first discovered in 1962 by Copp and Cheney [109]. The sequencing of the porcine calcitonin in 1968 revealed that this polypeptide is composed of 32 amino acids [110]. In vertebrates, calcitonin is produced by the thyroid gland [111] and regulates calcium (Ca^2+^) levels in the blood, antagonizing the effects of parathyroid hormone [112, 113]. The evolutionary antiquity of calcitonin-related peptides was first revealed with the discovery that a diuretic hormone in insects (DH_31_) is a calcitonin-like peptide [114]. However, DH_31_ shares modest sequence similarity with vertebrate calcitonins and lacks the N-terminal disulphide bridge that is characteristic of calcitonin-type peptides in vertebrates. More recently, it has been discovered that both DH_31_-type and vertebrate calcitonin-type neuropeptides occur in some protostomian invertebrates, including the annelid *Platynereis dumerilii* and the insect *Locusta migratoria* [4, 115]. Hence, it is proposed that an ancestral-type calcitonin precursor gene duplicated in the common ancestor of protostomes to give rise to DH_31_-type and calcitonin-type peptides, but with subsequent loss of calcitonin-type peptides in some protostomes. Consistent with this hypothesis, calcitonin-type precursors but not DH_31_-type precursors have been identified in deuterostomian invertebrates, including echinoderms [8, 15, 16, 116].

An interesting feature of calcitonin/DH_31_ precursors is the occurrence of multiple splice variants. In vertebrates, alternative splicing of the calcitonin gene results in two transcripts: one transcript encodes calcitonin and the other transcript encodes calcitonin gene-related peptide [117]. Furthermore, a complex interplay of receptors and accessory proteins determines the pharmacological profile of these peptides [118, 119]. Alternative splicing of DH_31_ and calcitonin precursors in insects has also been previously reported [115, 120, 121]. Interestingly, alternative splicing of insect calcitonin genes also generates variants that give rise to different mature peptides [115]. However, unlike the calcitonin gene, DH_31_ splice variants all produce an identical mature peptide [120, 121].

Our analysis of the ophiuroid transcriptomes also identified two transcript variants for calcitonin (**Figure 7E and F**). Based on our analysis of transcript sequences, ophiuroid calcitonin genes comprise at least three putative coding regions or ‘exons’. It is unclear if these three coding regions represent three or more exons due to the lack of genomic data, but for the sake of simplicity, we refer to them here as ‘exons’. Transcript variant 1 comprises ‘exons’ 1 and 3 but lacks ‘exon’ 2 whereas transcript variant 2 contains all 3 ‘exons’. Interestingly, ‘exons’ 2 and 3 both encode a calcitonin-type peptide. Hence, transcript variant 1 encodes a precursor that produces one calcitonin-type peptide and transcript variant 2 encodes two non-identical calcitonin-type peptides. These alternatively spliced transcripts were found in several brittle star species (**Figure 8**) and thus this may represent an ancient and conserved feature, although transcript variant 1 was not found in *O. victoriae*.

**Figure 8.**
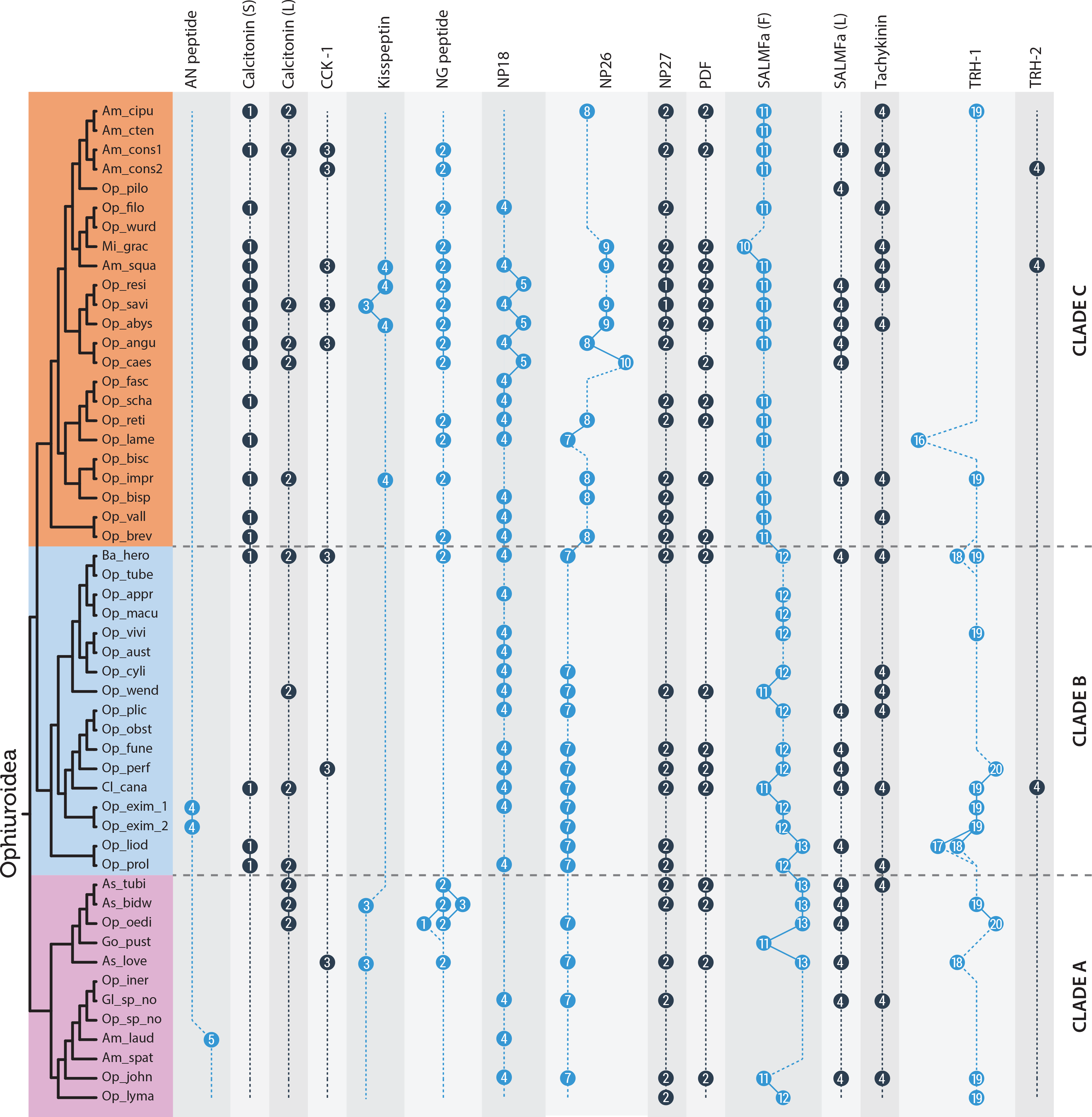
Comparison of neuropeptide copy numbers across the Ophiuroidea for precursors comprising multiple copies of neuropeptides. Neuropeptide precursors were mined from 52 ophiuroid transcriptomes, with the phylogeny adapted from O’Hara et al. (2014) [12]. Am_laud: *Amphiophiura laudata*, Am_spat: *Amphiophiura spatulifera*, Am_cipu: *Amphioplus cipus*, Am_cten: *Amphioplus ctenacantha*, Am_squa: *Amphipholis squamata*, Am_cons1: *Amphiura constricta* 1, Am_cons2: *Amphiura constricta* 2, As_love: *Asteronyx loveni*, As_bidw: *Asteroschema bidwillae*, As_tubi: *Asteroschema tubiferum*, Ba_hero: *Bathypectinura heros*, Cl_cana: *Clarkcoma canaliculata*, Gl_sp_no: *Glaciacantha sp* nov, Go_pust: *Gorgonocephalus pustulatum*, Mi_grac: *Microphiopholis gracillima*, Op_fune: *Ophiacantha funebris*, Op_abys: *Ophiactis abyssicola*, Op_resi: *Ophiactis resiliens*, Op_savi: *Ophiactis savignyi*, Op_vall: *Ophiernus vallincola*, Op_pilo: *Ophiocentrus pilosus*, Op_wend: *Ophiocoma wendtii*, Op_oedi: *Ophiocreas oedipus*, Op_tube: *Ophiocypris tuberculosis*, Op_appr: *Ophioderma appressum*, Op_bisc: *Ophiolepis biscalata*, Op_impr: *Ophiolepis impressa*, Op_brev: *Ophioleuce brevispinum*, Op_perf: *Ophiolimna perfida*, Op_prol: *Ophiologimus prolifer*, Op_obst: *Ophiomoeris obstricta*, Op_lyma: *Ophiomusium lymani*, Op_aust: *Ophiomyxa australis*, Op_vivi: *Ophiomyxa* sp cf *vivipara*, Op_fasc: *Ophionereis fasciata*, Op_reti: *Ophionereis reticulata*, Op_scha: *Ophionereis schayeri*, Op_cyli: *Ophiopeza cylindrica*, Op_filo: *Ophiophragmus filograneus*, Op_wurd: *Ophiophragmus wurdemanii*, Op_liod: *Ophiophrura liodisca*, Op_john: *Ophiophycis johni*, Op_lame: *Ophioplax lamellosa*, Op_iner: *Ophiopleura inermis*, Op_plic: *Ophioplinthaca plicata*, Op_bisp: *Ophioplocus bispinosus*, Op_macu: *Ophiopsammus maculata*, Op_angu: *Ophiothrix angulata*, Op_caes: *Ophiothrix caespitosa*, Op_exim_1: *Ophiotreta eximia* 1, Op_exim_2: *Ophiotreta eximia* 2, Op_sp_no: *Ophiura sp nov*.

Previous studies have identified precursors comprising a single calcitonin-type peptide in the starfish *A. rubens* and the sea urchin *S. purpuratus* [8, 16], and a precursor comprising two calcitonin-type peptides in the sea cucumber *A. japonicus* [15]. Informed by the identification here of two transcript types in ophiuroids (transcript variant 1 and 2), we have now discovered that two transcript types also occur in *A. japonicus* transcriptome. Hence, alternative splicing of calcitonin-type precursor genes can be traced back in the echinoderm lineage to the common ancestor of the Asterozoa and Echinozoa, but with subsequent loss of this characteristic in some lineages.

#### GPA2 and GPB5

The vertebrate glycoprotein hormone family comprises luteinizing hormone (LH) follicle-stimulating hormone (FSH), chorionic gonadotropin (CG), thyroid-stimulating hormone (TSH) and the recently discovered thyrostimulin (TS) [122, 123]. Thyrostimulin is a heterodimer composed of two subunits, glycoprotein alpha 2 (GPA2) and glycoprotein beta 5 (GPB5). Orthologs of GPA2 and GPB5 have been identified and characterized in the insect *Drosophila melanogaster* [124] and in other invertebrates, including echinoderms [125]. Insect GPA2 and GPB5 both contain 10 conserved cysteine residues that are important in forming a heterodimeric cysteine-knot structure. Surprisingly, *A. japonicus* GPA2 contains only 7 cysteine residues (having lost residues 7, 8 and 9) while *O. victoriae* GPB5.1, *A. rubens* GPB5.1 and *S. purpuratus* GPB5 all contain 8 cysteine residues (having lost the final two cysteine residues) (**Figure S5**). It is difficult to predict the structural differences that may arise in the heterodimer due to this variability in the number of cysteine residues. The possibility of GPA2 and/or GPB5 monomers or homodimers exerting their own biological functions has not been ruled out [126]. Additional investigations are needed to investigate if GPA2 and GPB5 are co-localized in echinoderms and if the monomers and dimers (both homo and hetero) exert different effects.

### Uncharacterized neuropeptides

In addition to the neuropeptides discussed above, we have also identified three neuropeptide precursors that could not be classified into any known neuropeptide families. These include *O. victoriae* neuropeptide precursor (Ovnp) 18 (*O. victoriae* ortholog of Spnp18 in *S. purpuratus*) [16], Ovnp26 and Ovnp27, with the latter two identified for the first time in echinoderms. The choice of nomenclature for Ovnp26 and Ovnp27 is based on a previously used numerical nomenclature in *S. purpuratus* and/or *A. rubens*, which goes up to Arnp25 in *A. rubens*.

#### Ovnp18

Ovnp18 comprises four copies of a predicted mature peptide with the sequence LFWVD and the C-terminal region of the precursor (partial sequence) contains at least four cysteine residues (**Figure 5F**, GenBank; MF155258). Interestingly, this precursor type only comprises a single mature peptide in *A. rubens, S. purpuratus* and *A. japonicus* and the C-terminal region contains 9, 8 and 8 cysteine residues, respectively (data not shown) [8, 15, 16].

#### Ovnp26

Ovnp26 was identified following an analysis of *O. victoriae* transcriptome sequence using NpSearch [8]. Orthologs of Ovnp26 (GenBank; MF155259) were identified in other brittle stars but not in other echinoderms (**Figure S2–S4**). Ovnp26 comprises seven copies of peptides with a conserved C-terminal GW motif, whereas orthologs in *O. aranea* and *A. filiformis* are predicted to generate eight copies of the mature peptide. Some of the mature peptides have a C-terminal SGW motif, which is similar to the C-terminus of predicted mature peptides derived from *O. victoriae* pedal peptide precursor 3 (**Figure S7**). However, the lack of sequence similarity in other parts of the peptide suggests that the C-terminal similarity may reflect convergence rather than homology.

#### Ovnp27

Ovnp27 (GenBank; MF155251) was identified following a HMM-based search for SIFamide-type peptides [127, 128], albeit with a high E-value. This neuropeptide precursor comprises two putative amidated mature peptides that are located immediately after the signal peptide (**Figure S2–S4**), as seen in SIFamide precursors [129]. The first peptide of the *O. victoriae* precursor has a C-terminal IFamide motif just like in insect SIFamides (**Figure S9**). However, there is no sequence similarity with SIFamides in the rest of the peptide. This coupled with the fact that SIFamide-type receptors have not been identified in echinoderms [6] suggests that the sequence similarity that peptides derived from Ovnp27-type precursors share with SIFamides may reflect convergence rather than homology.

### Neuropeptide precursors not found in brittle stars

Our analysis of ophiuroid transcriptome sequence data did not reveal orthologs of the Spnp9 precursor from *S. purpuratus* or the Arnp21, Arnp22, Arnp23 and Arnp24 precursors from *A. rubens* [8, 16]. An Spnp9 ortholog is found in *A. japonicus* but not in *A. rubens* [15] and therefore this neuropeptide precursor type may be restricted to the Echinozoa. Orthologs of Arnp21-24 have not been found in *O. victoriae, S. purpuratus* or *A. japonicus*, which suggests that these may be Asteroidea-specific precursors.

Previous studies have shown that receptors for leucokinin, ecdysis-triggering hormone, QRFP, parathyroid hormone, galanin/allatostatins-A and Neuromedin-U/CAPA are present in ambulacraria [6, 7, 15]. The presence of these receptors suggests that their cognate ligands should also be present in ambulacraria. However, our search approaches failed to identify any proteins in ophiuroids that resemble precursors of these neuropeptides.

### Evolutionary conservation and variation of neuropeptide copy number in the Ophiuroidea

Many neuropeptide precursors comprise several structurally similar but non-identical bioactive peptides–i.e. the precursor protein gives rise to a neuropeptide “cocktail”. This feature of neuropeptide precursors occurs throughout metazoans. But how do these “cocktails” of neuropeptides evolve and what is their functional significance? Are the copies of mature peptides functionally redundant or do they have their own specific functions? These are important questions in neuroendocrinology for which answers remain elusive.

Evidence that neuropeptide copy number may be functionally important has been obtained from comparison of the sequences of neuropeptide precursors in twelve *Drosophila* species, the common ancestor of which dates back ~50 million years [130]. The number of peptide copies in each neuropeptide precursor was found to be identical (except for the FMRFamide precursor) when compared between the twelve species, suggesting that stabilising selection has acted to conserve neuropeptide “cocktails” in the *Drosophila* lineage.

Here, a comparison of *O. victoriae, A. filiformis* and *O. aranea* neuropeptide precursors and their putative mature peptides revealed that fourteen neuropeptide precursors comprised multiple neuropeptide copies. In certain cases, the number of the mature peptides derived from a particular precursor varied across species, whereas in other cases the numbers remained constant (**Figure 4**). Interestingly, these three species belong to two of the three major clades of brittle stars that evolved ~270 million years ago [12]. While *O. victoriae* belongs to the Chilophiurina infraorder (clade A), *A. filiformis* and *O. aranea* belong to the Gnathophiurina infraorder (clade C). Hence, this prompted us to examine the evolution of neuropeptides and neuropeptide copy number variation at a higher level of phylogenetic resolution. To do this, we utilized a unique dataset comprising 52 ophiuroid transcriptomes. These transcriptomes were recently used as part of a phylotranscriptomic approach to reconstruct the phylogeny of ophiuroids, generating a robust phylogenetic tree that comprises three major clades [12]. Hence, this dataset allowed us to explore the evolution of neuropeptide precursors in the context of an established phylogenetic framework spanning over an unprecedented timescale of ~270 million years.

We selected for analysis neuropeptide precursors comprising more than a one putative mature neuropeptide, which include AN peptide, calcitonin, cholecystokinin 1, kisspeptin, np18, np26, np27, NG peptide, PDF, SALMFamide (L-type and F-type), tachykinin and TRH (1 and 2). Pedal peptide precursors (1, 2 and 3) were excluded from the analysis because orthology relationships between these precursors could not be established with confidence across all species (data not shown). We used *O. victoriae* representatives of these neuropeptide precursor families and the *A. filiformis* AN peptide precursor to mine 52 ophiuroid transcriptomes using BLAST. Multiple sequence alignments were generated based on the search hits (**Figure S11**) and the number of predicted mature peptides were compared (**Figure 8**). Interestingly, the number of peptides within the majority of precursors remained constant across all the species examined, which share a common ancestor estimated to date from ~270 million years ago [12].

Some studies that have investigated the physiological significance of neuropeptide “cocktails” indicate that neuropeptides derived from the same precursor protein are functionally redundant. For example, this was found for myomodulin neuropeptides in the mollusk *Aplysia californica* using the accessory radula closer muscle preparation as a bioassay [131] and for FMRFamide-related neuropeptides in *Drosophila melanogaster* when analysing effects on nerve-stimulated contraction of larval body-wall muscles [132]. However, the authors of the latter study cautiously highlighted the need to “search for additional functions or processes in which these peptides may act differentially”. Importantly, studies employing use of multiple bioassays have obtained data indicating that neuropeptides derived from a single precursor protein are not functionally redundant. For example, when the actions of fourteen structurally related neuropeptides derived from a precursor of *Mytilus* Inhibitory Peptide-related peptides in *Aplysia* were tested on three organ preparations (oesophagus, penis retractor, body wall) it was found that the rank order of potency for the peptides differed between preparations [133]. Similarly, when assaying the effects of allatostatin neuropeptides in cockroaches, tissue-specific differences in potency were observed [134]. The conservation of peptide copy number across a timescale of ~270 million years in the Ophiuroidea supports the idea that the occurrence of multiple copies of identical or structurally related neuropeptides is functionally important.

For those neuropeptide precursors that did exhibit variation in neuropeptide copy number, TRH-type precursors exhibited the highest variation, with numbers ranging from 16 to 20 copies (**Figure 9**). F-type SALMFamide precusors also showed variation in copy numbers (**Figure 10**) but loss of peptides was more frequent in F-type SALMFamide precursors than in TRH-type precursors. Furthermore, detailed analysis of sequence alignments for these precursors revealed that loss of neuropeptide copies is usually a consequence of non-synonymous mutations in codons for residues that form dibasic cleavage sites or for glycine residues that are substrates for the C-terminal amidation. This is not surprising since the C-terminal amide in smaller-sized peptides is usually important for receptor binding and activation. What is unclear at the moment is how the peptide copy number increases within a given precursor. Perhaps the increase in peptide copy number occurs as a result of unequal crossing-over during recombination [130].

**Figure 9.**
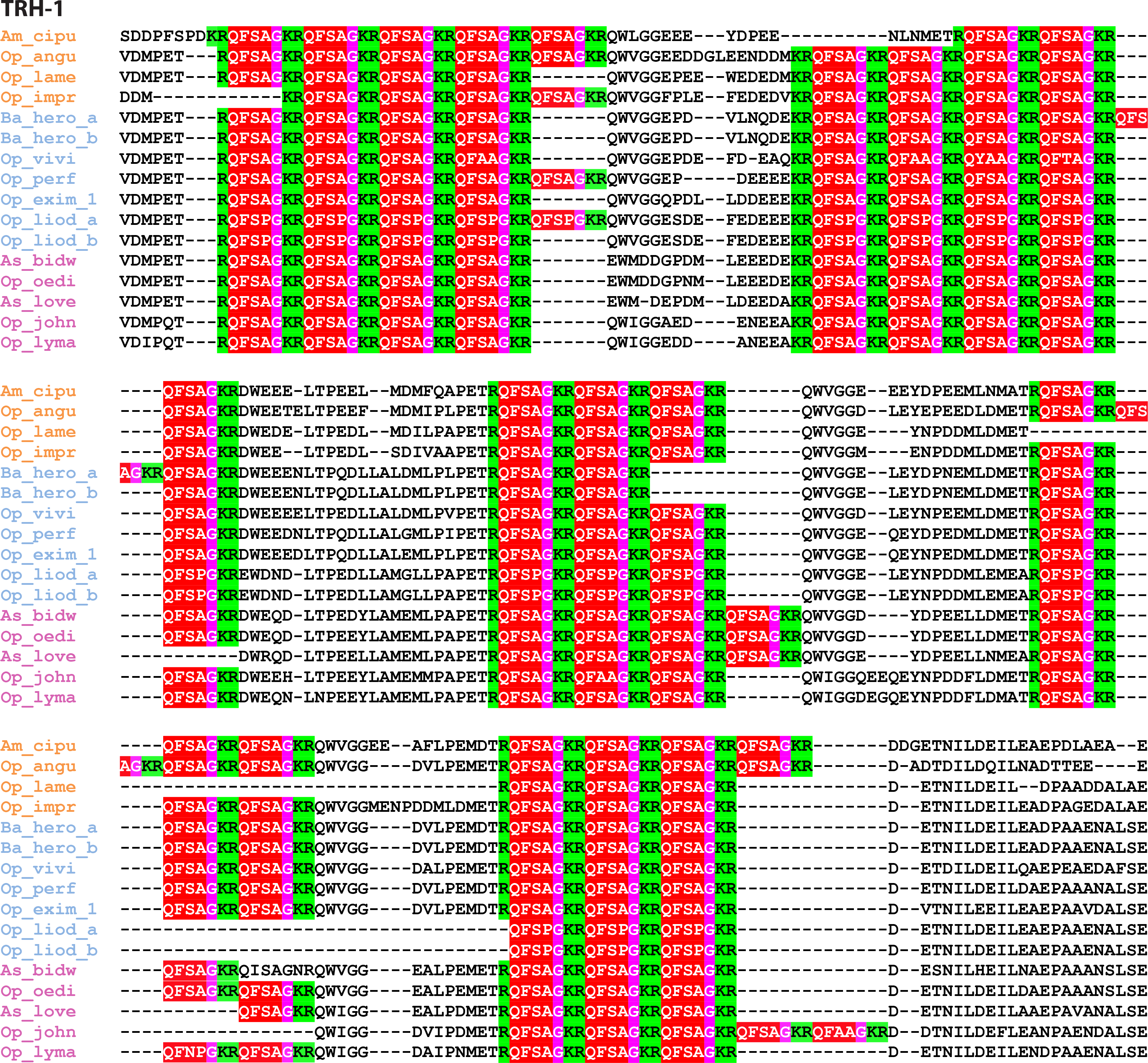
A partial multiple sequence alignment of ophiuroid thyrotropin-releasing hormone (TRH) precursors showing clade-specific gain/loss of neuropeptide copies. Mono- and di-basic cleavage sites are highlighted in green, mature peptides in red with the glycine residue for amidation in pink. Species have been grouped and coloured (clade A in purple, clade B in blue and clade C in orange) based on the phylogeny determined by O’Hara et al. (2014) [12].

**Figure 10.**
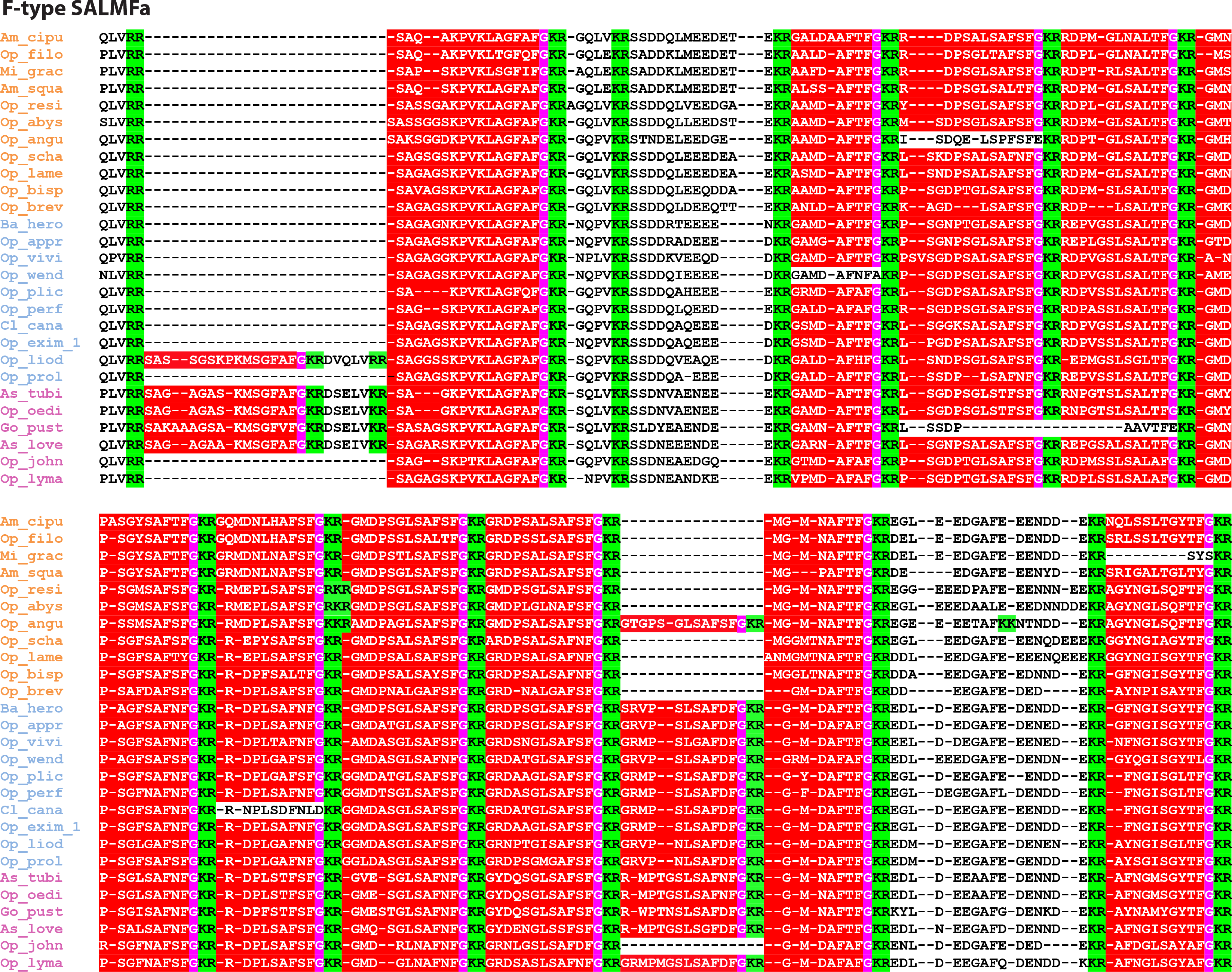
A partial multiple sequence alignment of ophiuroid F-type SALMFamide precursors showing clade-specific gain/loss of neuropeptide copies. Di-basic cleavage sites are highlighted in green, mature peptides in red with the glycine residue for amidation in pink. Species have been grouped and coloured (clade A in purple, clade B in blue and clade C in orange) based on the phylogeny determined by O’Hara et al. (2014) [12].

The number of peptides within the F-type SALMFamide precursors appear to be clade specific. Thus, the average/median number of F-type SALMFamides in precursors from clade A is 13, clade B is 12 and clade C is 11, with a few exceptions (**Figure 8**). Similarly, the number of peptides within NP26-type precursors also appears to be clade specific. Hence the number of peptides is highly stable at 7 peptides within clades A and B but a high variation in peptide copy number is observed in clade C. When examining peptide copy number within clades, there are a few cases where the number of peptides within a given precursor for certain species appears to be an exception/outlier. For instance, 16 copies of the mature peptide in *Ophioplax lamellosa* TRH-1 precursor is distinctly different to the 19 copies found in other species within that clade (clade C). Likewise, *Ophiactis savignyi* only has 3 copies of kisspeptin-type peptides compared to 4 copies found in other species of that clade (**Figure 8**).

It could be argued that misalignments during transcriptome assembly may have influenced the number of predicted peptides found in a given precursor. However, it is unlikely that misalignments have affected the predicted sequences of neuropeptide precursors comprising multiple copies of peptides that are similar but non-identical, which applies to the majority of the precursor proteins analysed here in ophiuroids. The only exception to this are the TRH-type precursors, where the encoded peptide sequences are short and often identical, even at the nucleotide level (data not shown), Another limitation of using transcriptome data is that the sequences of neuropeptide precursors may be partial or unknown for some species and where this applies a peptide copy number is not shown in Fig. 8. An extreme example of this is the AN peptide precursor, where complete precursors sequences were only obtained from the three reference species and three other species. However, for the majority of precursor types, sequence data was obtained from a variety of species from each of the three clades of ophiuroids. For example, complete F-type SALMFamide precursor sequences were found in most of the investigated species (39 species + 3 reference species).

## Conclusion

Here we report the first detailed analysis of the neuropeptide precursor complement of ophiuroids and the most comprehensive identification of echinoderm neuropeptide precursors to date. We have identified novel representatives of several bilaterian neuropeptide families in echinoderms for the first time, which include orthologs of endothelin/CCHamide, eclosion hormone, neuropeptide-F/Y and nucleobinin/nesfatin. Furthermore, analysis of precursor proteins comprising multiple copies of identical or related neuropeptides across ~270 million years of ophiuroid evolution indicates that the precise composition of neuropeptide “cocktails” is functionally important as evident from the conservation of neuropeptide copy number for multiple precursors.

## Methods

### Sequencing and assembly of transcriptomes

Ophiuroid transcriptomes used in this study were sequenced and assembled as reported previously [12, 20, 24].

### Identification of neuropeptide precursors in ophiuroids

In order to identify neuropeptide precursors in *O. victoriae, A. filiformis* and *O. aranea*, sequences of neuropeptide precursors identified previously in other echinoderms (including the starfish, *A. rubens*, the sea urchin *S. purpuratus* and the sea cucumber, *A. japonicus*) were used as queries for tBLASTn analysis of a transcriptome database, using an e value of 1000. Sequences identified as potential neuropeptide precursors by BLAST were translated using the ExPASy Translate tool (http://web.expasy.org/translate/) and then analysed for features of neuropeptide precursors. Specifically, sequences were evaluated based on 1) the presence of an N-terminal signal peptide (using Signal P v 4.1 with the sensitive cut-off of 0.34) and 2) the presence of monobasic or dibasic cleavage sites flanking the putative bioactive peptide(s).

To identify novel neuropeptide precursors or highly-divergent precursors with low sequence similarity to known precursors, we utilized two additional approaches. In the first approach, we used NpSearch [8], software that identifies putative neuropeptide precursors based on various characteristics (presence of signal peptide and dibasic cleavage sites amongst others). In the second approach, NpHMMer (http://nphmmer.sbcs.qmul.ac.uk/), a Hidden Markov Models (HMM) based software was used to identify neuropeptides not found using the above approaches.

Neuropeptide precursors identified in *O. victoriae* (which represented a more comprehensive neuropeptide precursor repertoire compared to *A. filiformis* and *O. aranea*) were then submitted as queries for BLAST analysis of sequence data from 52 Ophiuroidea species, using an E-value of 1e-06. BLAST hits were then further analysed using an automated ruby script (available at https://github.com/IsmailM/ophiuroid_neuropeptidome). Each BLAST hit was translated using BioRuby and the open reading frame (ORF) containing the BLAST high-scoring segment pair was extracted. These ORFs were then examined for the presence of a signal peptide using Signal P 4.1 using a sensitive cut-off of 0.34. All sequences were then aligned using MAFFT, with the number of maximum iterations set to 1000 to ensure an optimal alignment. These alignments were then further optimized by manually adjusting the location of the bioactive peptide and cleavage sites. Finally, the alignments were annotated using different colours for the signal peptide (blue), the bioactive peptide(s) (red) and cleavage sites (green).

### Phylogenetic and clustering analyses of sequence data

Phylogenetic analysis of membrane guanylyl cyclase receptors and nucleobindins was performed using maximuim likelihood and Bayesian methods. Prior to these analyses, corresponding multiple alignments were trimmed using BMGE [135] with the following options: BLOSUM30, max–h = 1, − b = 1, as described previously [10, 94]. The maximum likelihood method was implemented in the PhyML program (v3.1/3.0 aLRT). The WAG substitution model was selected assuming an estimated proportion of invariant sites (of 0.112) and 4 gamma-distributed rate categories to account for rate heterogeneity across sites. The gamma shape parameter was estimated directly from the data. Reliability for internal branch was assessed using the bootstrapping method (500 bootstrap replicates). The Bayesian inference method was implemented in the MrBayes program (v3.2.3). The number of substitution types was fixed to 6. The poisson model was used for substitution, while rates variation across sites was fixed to “invgamma”. Four Markov Chain Monte Carlo (MCMC) chains were run for 100000 generations, sampling every 100 generations, with the first 500 sampled trees discarded as “burn-in”. Finally, a 50% majority rule consensus tree was constructed.

CLANS analysis was performed on echinoderm EH-like, arthropod EH, arthropod ITP and vertebrates ANP precursors based on all-against-all sequence similarity (BLAST searches) using BLOSUM 45 matrix (https://toolkit.tuebingen.mpg.de/clans/) [41] and the significant high-scoring segment pairs (HSPs). Neuropeptide precursors were clustered in a three-dimensional graph represented here in two dimensions.

## Data accessibility

Raw sequence data used to assemble the transcriptomes have been deposited in the NCBI Sequence Read Archive (SRA) under the accession number SRP107914 (https://www.ncbi.nlm.nih.gov/sra/?term=SRP107914) and in the NCBI BioProject under the accession number PRJNA311384 (https://www.ncbi.nlm.nih.gov/bioproject/311384).

## Author contributions

M.Z., T.D.O. and M.R.E.: designed the research; I.M.: generated HMM models; M.Z., I.M., L.A.Y.G., J.D., N.A. and A.F.H: identified the neuropeptide precursors; M.Z., I.M., L.A.Y.G., J.D. and N.A.: analysed the data; M.Z., J.D. and M.R.E. wrote the manuscript with input from other authors. M.Z. and M.R.E: supervised the study.

## Acknowledgements

The authors would like to acknowledge Zuraiha Waffa and Giulia Oluwabunmi Olayemi for their assistance with sequence alignments.

## Supplementary files

**Figure S1.** Alignment and phylogenetic analysis of nucleobindins (NUCB). A) Partial sequence alignment (excludes the signal peptide) of NUCB precursors. The locations of *Homo sapiens* nesfatin-1, 2 and 3 are indicated. A dibasic cleavage site in *O. victoriae* nesfatin-1 is marked in red. B) Phylogenetic analysis of NUCB precursors. Species names: *Ophionotus victoriae* (Ovic), *Amphiura filiformis* (Afil), *Ophiopsila aranea* (Oara), *Apostichopus japonicus* (Ajap), *Strongylocentrotus purpuratus* (Spur), *Homo sapiens* (Hsap), *Mus musculus* (Mmus) and *Drosophila melanogaster* (Dmel).

**Figure S2.** *Ophionotus victoriae* neuropeptide precursor repertoire.

**Figure S3.** *Amphiura filiformis* neuropeptide precursor repertoire.

**Figure S4.** *Ophiopsila aranea* neuropeptide precursor repertoire.

**Figure S5.** Partial multiple sequence alignments of echinoderm representatives of A) glycoprotein alpha 2 (GPA2)-type subunits and B) glycoprotein beta 5 (GPB5)-type subunits. Species names: *Ophionotus victoriae* (Ovic), *Asterias rubens* (Arub), *Strongylocentrotus purpuratus* (Spur) and *Apostichopus japonicus* (Ajap).

**Figure S6.** Partial multiple sequence alignments of echinoderm representatives of large protein hormones. A) insulin/insulin-like growth factor; B) relaxin-like peptide; C) bursicon (bursicon alpha); D) partner of bursicon (bursicon beta). Species names: *Ophionotus victoriae* (Ovic), *Asterias rubens* (Arub), *Strongylocentrotus purpuratus* (Spur) and *Apostichopus japonicus* (Ajap).

**Figure S7.** Multipe sequence alignment of echinoderm pedal peptides. Species names: *Ophionotus victoriae* (Ovic), *Asterias rubens* (Arub), *Strongylocentrotus purpuratus* (Spur) and *Apostichopus japonicus* (Ajap).

**Figure S8.** Multiple sequence alignments of echinoderm neuropeptide families. A) F-type SALMFamide alignment; B) L-type SALMFamide alignment; C) AN peptide. Species names: *Ophionotus victoriae* (Ovic), *Asterias rubens* (Arub), *Strongylocentrotus purpuratus* (Spur) and *Apostichopus japonicus* (Ajap).

**Figure S9.** Multiple sequence alignment of predicted peptides derived from neuropeptide precursor 27 in *Ophionotus victoriae* (Ovic), *Amphiura filiformis* (Afil), *Ophiopsila aranea* (Oara) and *Apostichopus japonicus* (Ajap).

**Figure S10.** Multiple sequence alignment of pigment-dispersing factor-type precursors. Note the conservation of cleavage sites (KR) immediately preceding the mature peptide as well as the location of the mature peptide (C-terminal end of the precursor). Species names: *Ophionotus victoriae* (Ovic), *Asterias rubens* (Arub), *Aplysia californica* (Acal), *Platynereis dumerilii* (Pdum), *Euperipatoides rowelli* (Erow), *Nilaparvata lugens* (Nlug), *Bombyx mori* (Bmor) and *Drosophila melanogaster* (Dmel).

**Figure S11.** Multiple sequence alignments of neuropeptide precursors used to generate Figure 8.

**Figure S12.** Partial nucleotide sequence of the *Ophionotus victoriae* neuropeptide Y/F precursor.

## References

1 Nassel, D. R., Winther, A. M. 2010 *Drosophila* neuropeptides in regulation of physiology and behavior. Prog Neurobiol. 92, 42–104. (10.1016/j.pneurobio.2010.04.010)

2 Argiolas, A., Melis, M. R. 2013 Neuropeptides and central control of sexual behaviour from the past to the present: a review. Prog Neurobiol. 108, 80–107. (10.1016/j.pneurobio.2013.06.006)

3 Sohn, J. W., Elmquist, J. K., Williams, K. W. 2013 Neuronal circuits that regulate feeding behavior and metabolism. Trends Neurosci. 36, 504–512. (10.1016/j.tins.2013.05.003)

4 Conzelmann, M., Williams, E. A., Krug, K., Franz-Wachtel, M., Macek, B., Jekely, G. 2013 The neuropeptide complement of the marine annelid Platynereis dumerilii. BMC Genomics. 14, 906. (10.1186/1471-2164-14-906)

5 Veenstra, J. A. 2016 Neuropeptide Evolution: Chelicerate Neurohormone and Neuropeptide Genes may reflect one or more whole genome duplications. Gen Comp Endocrinol. (10.1016/j.ygcen.2015.07.014)

6 Mirabeau, O., Joly, J. S. 2013 Molecular evolution of peptidergic signaling systems in bilaterians. Proc Natl Acad Sci U S A. 110, E2028–2037. (10.1073/pnas.1219956110)

7 Jekely, G. 2013 Global view of the evolution and diversity of metazoan neuropeptide signaling. Proc Natl Acad Sci U S A. 110, 8702–8707. (10.1073/pnas.1221833110)

8 Semmens, D. C., Mirabeau, O., Moghul, I., Pancholi, M. R., Wurm, Y., Elphick, M. R. 2016 Transcriptomic identification of starfish neuropeptide precursors yields new insights into neuropeptide evolution. Open Biol. 6, 150224. (10.1098/rsob.150224)

9 Semmens, D. C., Beets, I., Rowe, M. L., Blowes, L. M., Oliveri, P., Elphick, M. R. 2015 Discovery of sea urchin NGFFFamide receptor unites a bilaterian neuropeptide family. Open Biol. 5, 150030. (10.1098/rsob.150030)

10 Tian, S., Zandawala, M., Beets, I., Baytemur, E., Slade, S. E., Scrivens, J. H., Elphick, M. R. 2016 Urbilaterian origin of paralogous GnRH and corazonin neuropeptide signalling pathways. Sci Rep. 6, 28788. (10.1038/srep28788)

11 Telford, M. J., Lowe, C. J., Cameron, C. B., Ortega-Martinez, O., Aronowicz, J., Oliveri, P., Copley, R. R. 2014 Phylogenomic analysis of echinoderm class relationships supports Asterozoa. Proc Biol Sci. 281, (10.1098/rspb.2014.0479)

12 O’Hara, T. D., Hugall, A. F., Thuy, B., Moussalli, A. 2014 Phylogenomic resolution of the class Ophiuroidea unlocks a global microfossil record. Current Biology. 24, 1874–1879.

13 Hyman, L. H. 1955 The invertebrates. Vol. 4. Echinodermata, 763 pp. MacGraw-Hill: New York.

14 Wilkie, I. 2001 Autotomy as a prelude to regeneration in echinoderms. Microscopy research and technique. 55, 369–396.

15 Rowe, M. L., Achhala, S., Elphick, M. R. 2014 Neuropeptides and polypeptide hormones in echinoderms: new insights from analysis of the transcriptome of the sea cucumber Apostichopus japonicus. Gen Comp Endocrinol. 197, 43–55. (10.1016/j.ygcen.2013.12.002)

16 Rowe, M. L., Elphick, M. R. 2012 The neuropeptide transcriptome of a model echinoderm, the sea urchin *Strongylocentrotus purpuratus*. Gen Comp Endocrinol. 179, 331–344. (10.1016/j.ygcen.2012.09.009)

17 Mayorova, T. D., Tian, S., Cai, W., Semmens, D. C., Odekunle, E. A., Zandawala, M., Badi, Y., Rowe, M. L., Egertova, M., Elphick, M. R. 2016 Localization of Neuropeptide Gene Expression in Larvae of an Echinoderm, the Starfish *Asterias rubens*. Front Neurosci. 10, 553. (10.3389/fnins.2016.00553)

18 Kim, C. H., Kim, E. J., Go, H. J., Oh, H. Y., Lin, M., Elphick, M. R., Park, N. G. 2016 Identification of a novel starfish neuropeptide that acts as a muscle relaxant. J Neurochem. 137, 33–45. (10.1111/jnc.13543)

19 Jones, C. E., Zandawala, M., Semmens, D. C., Anderson, S., Hanson, G. R., Janies, D. A., Elphick, M. R. 2016 Identification of a neuropeptide precursor protein that gives rise to a “cocktail” of peptides that bind Cu(II) and generate metal-linked dimers. Biochim Biophys Acta. 1860, 57–66. (10.1016/j.bbagen.2015.10.008)

20 Elphick, M. R., Semmens, D. C., Blowes, L. M., Levine, J., Lowe, C. J., Arnone, M. I., Clark, M. S. 2015 Reconstructing SALMFamide Neuropeptide Precursor Evolution in the Phylum Echinodermata: Ophiuroid and Crinoid Sequence Data Provide New Insights. Front Endocrinol (Lausanne). 6, 2. (10.3389/fendo.2015.00002)

21 Lin, M., Mita, M., Egertova, M., Zampronio, C. G., Jones, A. M., Elphick, M. R. 2016 Cellular localization of relaxin-like gonad-stimulating peptide expression in Asterias rubens: New insights into neurohormonal control of spawning in starfish. J Comp Neurol. (10.1002/cne.24141)

22 Stöhr, S., O’Hara, T. D., Thuy, B. 2012 Global diversity of brittle stars (Echinodermata: Ophiuroidea). PLoS One. 7, e31940.

23 Shackleton, J. D. 2005 Skeletal homologies, phylogeny and classification of the earliest asterozoan echinoderms. Journal of Systematic Palaeontology. 3, 29–114.

24 Delroisse, J., Mallefet, J., Flammang, P. 2016 De Novo Adult Transcriptomes of Two European Brittle Stars: Spotlight on Opsin-Based Photoreception. PLoS One. 11, e0152988. (10.1371/journal.pone.0152988)

25 Delroisse, J., Ortega-Martinez, O., Dupont, S., Mallefet, J., Flammang, P. 2015 De novo transcriptome of the European brittle star Amphiura filiformis pluteus larvae. Marine genomics. 23, 109–121.

26 Burns, G., Thorndyke, M. C., Peck, L. S., Clark, M. S. 2013 Transcriptome pyrosequencing of the Antarctic brittle star *Ophionotus victoriae*. Mar Genomics. 9, 9–15. (10.1016/j.margen.2012.05.003)

27 Cannon, J. T., Kocot, K. M., Waits, D. S., Weese, D. A., Swalla, B. J., Santos, S. R., Halanych, K. M. 2014 Phylogenomic resolution of the hemichordate and echinoderm clade. Curr Biol. 24, 2827–2832. (10.1016/j.cub.2014.10.016)

28 Vaughn, R., Garnhart, N., Garey, J. R., Thomas, W. K., Livingston, B. T. 2012 Sequencing and analysis of the gastrula transcriptome of the brittle star *Ophiocoma wendtii*. Evodevo. 3, 19. (10.1186/2041-9139-3-19)

29 Purushothaman, S., Saxena, S., Meghah, V., Swamy, C. V., Ortega-Martinez, O., Dupont, S., Idris, M. 2015 Transcriptomic and proteomic analyses of *Amphiura filiformis* arm tissue-undergoing regeneration. J Proteomics. 112, 113–124. (10.1016/j.jprot.2014.08.011)

30 Smith, A. B., Paterson, G. L., Lafay, B. 1995 Ophiuroid phylogeny and higher taxonomy: morphological, molecular and palaeontological perspectives. Zoological Journal of the Linnean Society. 114, 213–243.

31 Thuy, B., Stohr, S. 2016 A New Morphological Phylogeny of the Ophiuroidea (Echinodermata) Accords with Molecular Evidence and Renders Microfossils Accessible for Cladistics. PLoS One. 11, e0156140. (10.1371/journal.pone.0156140)

32 O’Hara, T. D., Hugall, A. F., Thuy, B., Stohr, S., Martynov, A. V. 2017 Restructuring higher taxonomy using broad-scale phylogenomics: The living Ophiuroidea. Mol Phylogenet Evol. 107, 415–430. (10.1016/j.ympev.2016.12.006)

33 Truman, J. W., Riddiford, L. M. 1970 Neuroendocrine control of ecdysis in silkmoths. Science. 167, 1624–1626. (10.1126/science.167.3925.1624)

34 Kataoka, H., Troetschler, R. G., Kramer, S. J., Cesarin, B. J., Schooley, D. A. 1987 Isolation and primary structure of the eclosion hormone of the tobacco hornworm, Manduca sexta. Biochem Biophys Res Commun. 146, 746–750.

35 Zhou, L., Li, S., Wang, Z., Li, F., Xiang, J. 2017 An eclosion hormone-like gene participates in the molting process of Palaemonid shrimp *Exopalaemon carinicauda*. Dev Genes Evol. 227, 189–199. (10.1007/s00427-017-0580-9)

36 Ewer, J. 2005 Behavioral actions of neuropeptides in invertebrates: insights from *Drosophila*. Horm Behav. 48, 418–429. (10.1016/j.yhbeh.2005.05.018)

37 Zitnan, D., Kim, Y. J., Zitnanova, I., Roller, L., Adams, M. E. 2007 Complex steroid-peptide-receptor cascade controls insect ecdysis. Gen Comp Endocrinol. 153, 88–96. (10.1016/j.ygcen.2007.04.002)

38 Chang, J. C., Yang, R. B., Adams, M. E., Lu, K. H. 2009 Receptor guanylyl cyclases in Inka cells targeted by eclosion hormone. Proc Natl Acad Sci U S A. 106, 13371–13376. (10.1073/pnas.0812593106)

39 McNabb, S. L., Baker, J. D., Agapite, J., Steller, H., Riddiford, L. M., Truman, J. W. 1997 Disruption of a behavioral sequence by targeted death of peptidergic neurons in Drosophila. Neuron. 19, 813–823.

40 Kruger, E., Mena, W., Lahr, E. C., Johnson, E. C., Ewer, J. 2015 Genetic analysis of Eclosion hormone action during *Drosophila* larval ecdysis. Development. 142, 4279–4287. (10.1242/dev.126995)

41 Frickey, T., Lupas, A. 2004 CLANS: a Java application for visualizing protein families based on pairwise similarity. Bioinformatics. 20, 3702–3704. (10.1093/bioinformatics/bth444)

42 Vogel, K. J., Brown, M. R., Strand, M. R. 2013 Phylogenetic investigation of Peptide hormone and growth factor receptors in five dipteran genomes. Front Endocrinol (Lausanne). 4, 193. (10.3389/fendo.2013.00193)

43 Roller, L., Yamanaka, N., Watanabe, K., Daubnerova, I., Zitnan, D., Kataoka, H., Tanaka, Y. 2008 The unique evolution of neuropeptide genes in the silkworm *Bombyx mori*. Insect Biochem Mol Biol. 38, 1147–1157.

44 Hansen, K. K., Hauser, F., Williamson, M., Weber, S. B., Grimmelikhuijzen, C. J. 2011 The Drosophila genes CG14593 and CG30106 code for G-protein-coupled receptors specifically activated by the neuropeptides CCHamide-1 and CCHamide-2. Biochem Biophys Res Commun. 404, 184–189. (10.1016/j.bbrc.2010.11.089)

45 Ida, T., Takahashi, T., Tominaga, H., Sato, T., Sano, H., Kume, K., Ozaki, M., Hiraguchi, T., Shiotani, H., Terajima, S., et al. 2012 Isolation of the bioactive peptides CCHamide-1 and CCHamide-2 from Drosophila and their putative role in appetite regulation as ligands for G protein-coupled receptors. Front Endocrinol (Lausanne). 3, 177. (10.3389/fendo.2012.00177)

46 Farhan, A., Gulati, J., Grobetae-Wilde, E., Vogel, H., Hansson, B. S., Knaden, M. 2013 The CCHamide 1 receptor modulates sensory perception and olfactory behavior in starved Drosophila. Sci Rep. 3, 2765. (10.1038/srep02765)

47 Ren, G. R., Hauser, F., Rewitz, K. F., Kondo, S., Engelbrecht, A. F., Didriksen, A. K., Schjott, S. R., Sembach, F. E., Li, S., Sogaard, K. C., et al. 2015 CCHamide-2 Is an Orexigenic Brain-Gut Peptide in Drosophila. PLoS One. 10, e0133017. (10.1371/journal.pone.0133017)

48 Veenstra, J. A. 2010 Neurohormones and neuropeptides encoded by the genome of *Lottia gigantea*, with reference to other mollusks and insects. Gen Comp Endocrinol. 167, 86–103. (10.1016/j.ygcen.2010.02.010)

49 Oumi, T., Ukena, K., Matsushima, O., Ikeda, T., Fujita, T., Minakata, H., Nomoto, K. 1995 The GGNG peptides: novel myoactive peptides isolated from the gut and the whole body of the earthworms. Biochem Biophys Res Commun. 216, 1072–1078. (10.1006/bbrc.1995.2730)

50 Tatemoto, K., Carlquist, M., Mutt, V. 1982 Neuropeptide Y–a novel brain peptide with structural similarities to peptide YY and pancreatic polypeptide. Nature. 296, 659–660.

51 Tatemoto, K. 1982 Neuropeptide Y: complete amino acid sequence of the brain peptide. Proc Natl Acad Sci U S A. 79, 5485–5489.

52 Nassel, D. R., Wegener, C. 2011 A comparative review of short and long neuropeptide F signaling in invertebrates: Any similarities to vertebrate neuropeptide Y signaling? Peptides. 32, 1335–1355. (10.1016/j.peptides.2011.03.013)

53 Reichmann, F., Holzer, P. 2016 Neuropeptide Y: A stressful review. Neuropeptides. 55, 99–109. (10.1016/j.npep.2015.09.008)

54 Haas, D. A., George, S. R. 1989 Neuropeptide Y-induced effects on hypothalamic corticotropin-releasing factor content and release are dependent on noradrenergic/adrenergic neurotransmission. Brain Res. 498, 333–338.

55 Maule, A. G., Shaw, C., Halton, D. W., Thim, L., Johnston, C. F., Fairweather, I., Buchanan, K. D. 1991 Neuropeptide-F-a Novel Parasitic Flatworm Regulatory Peptide from Moniezia-Expansa (Cestoda, Cyclophyllidea). Parasitology. 102, 309–316.

56 Elphick, M. R., Mirabeau, O. 2014 The Evolution and Variety of RFamide-Type Neuropeptides: Insights from Deuterostomian Invertebrates. Front Endocrinol (Lausanne). 5, 93. (10.3389/fendo.2014.00093)

57 Kanai, Y., Tanuma, S. 1992 Purification of a novel B cell growth and differentiation factor associated with lupus syndrome. Immunol Lett. 32, 43–48.

58 Miura, K., Titani, K., Kurosawa, Y., Kanai, Y. 1992 Molecular cloning of nucleobindin, a novel DNA-binding protein that contains both a signal peptide and a leucine zipper structure. Biochem Biophys Res Commun. 187, 375–380.

59 Gonzalez, R., Mohan, H., Unniappan, S. 2012 Nucleobindins: bioactive precursor proteins encoding putative endocrine factors? Gen Comp Endocrinol. 176, 341–346. (10.1016/j.ygcen.2011.11.021)

60 Oh, I. S., Shimizu, H., Satoh, T., Okada, S., Adachi, S., Inoue, K., Eguchi, H., Yamamoto, M., Imaki, T., Hashimoto, K., et al. 2006 Identification of nesfatin-1 as a satiety molecule in the hypothalamus. Nature. 443, 709–712. (10.1038/nature05162)

61 Dore, R., Levata, L., Lehnert, H., Schulz, C. 2017 Nesfatin-1: functions and physiology of a novel regulatory peptide. J Endocrinol. 232, R45–R65. (10.1530/JOE-16-0361)

62 Sundarrajan, L., Blanco, A. M., Bertucci, J. I., Ramesh, N., Canosa, L. F., Unniappan, S. 2016 Nesfatin-1-Like Peptide Encoded in Nucleobindin-1 in Goldfish is a Novel Anorexigen Modulated by Sex Steroids, Macronutrients and Daily Rhythm. Sci Rep. 6, 28377. (10.1038/srep28377)

63 Otte, S., Barnikol-Watanabe, S., Vorbruggen, G., Hilschmann, N. 1999 NUCB1, the *Drosophila melanogaster* homolog of the mammalian EF-hand proteins NEFA and nucleobindin. Mech Dev. 86, 155–158.

64 Burke, R. D., Angerer, L. M., Elphick, M. R., Humphrey, G. W., Yaguchi, S., Kiyama, T., Liang, S., Mu, X., Agca, C., Klein, W. H., et al. 2006 A genomic view of the sea urchin nervous system. Dev Biol. 300, 434–460. (10.1016/j.ydbio.2006.08.007)

65 Elphick, M. R., Rowe, M. L. 2009 NGFFFamide and echinotocin: structurally unrelated myoactive neuropeptides derived from neurophysin-containing precursors in sea urchins. J Exp Biol. 212, 1067–1077. (10.1242/jeb.027599)

66 Mita, M., Yoshikuni, M., Ohno, K., Shibata, Y., Paul-Prasanth, B., Pitchayawasin, S., Isobe, M., Nagahama, Y. 2009 A relaxin-like peptide purified from radial nerves induces oocyte maturation and ovulation in the starfish, Asterina pectinifera. Proc Natl Acad Sci U S A. 106, 9507–9512. (10.1073/pnas.0900243106)

67 Elphick, M. R., Reeve, J. R., Jr., Burke, R. D., Thorndyke, M. C. 1991 Isolation of the neuropeptide SALMFamide-1 from starfish using a new antiserum. Peptides. 12, 455–459.

68 Schally, A. V., Bowers, C. Y., Redding, T. W., Barrett, J. F. 1966 Isolation of thyrotropin releasing factor (TRF) from porcine hypothalamus. Biochem Biophys Res Commun. 25, 165–169.

69 Boler, J., Enzmann, F., Folkers, K., Bowers, C. Y., Schally, A. V. 1969 The identity of chemical and hormonal properties of the thyrotropin releasing hormone and pyroglutamyl-histidyl-proline amide. Biochem Biophys Res Commun. 37, 705–710.

70 Burgus, R., Dunn, T. F., Desiderio, D., Guillemin, R. 1969 [Molecular structure of the hypothalamic hypophysiotropic TRF factor of ovine origin: mass spectrometry demonstration of the PCA-His-Pro-NH2 sequence]. C R Acad Sci Hebd Seances Acad Sci D. 269, 1870–1873.

71 Guillemin, R., Yamazaki, E., Gard, D. A., Jutisz, M., Sakiz, E. 1963 *In Vitro* Secretion of Thyrotropin (TSH): Stimulation by a Hypothalamic Peptide (TRF). Endocrinology. 73, 564–572. (10.1210/endo-73-5-564)

72 Bowers, C. R., Redding, T. W., Schally, A. V. 1965 Effect of thyrotropin releasing factor (TRF) of ovine, bovine, porcine and human origin on thyrotropin release *in vitro* and *in vivo*. Endocrinology. 77, 609–616. (10.1210/endo-77-4-609)

73 Bauknecht, P., Jekely, G. 2015 Large-Scale Combinatorial Deorphanization of *Platynereis* Neuropeptide GPCRs. Cell Rep. 12, 684–693. (10.1016/j.celrep.2015.06.052)

74 Wilber, J. F., Yamada, M., Kim, U. J., Feng, P., Carnell, N. E. 1992 The human prepro thyrotropin-releasing hormone (TRH) gene: cloning, characterization, hormonal regulation, and gene localization. Trans Am Clin Climatol Assoc. 103, 111–119.

75 Mutt, V., Jorpes, J. E. 1968 Structure of Porcine Cholecystokinin-Pancreozymin. I. Cleavage with Thrombin and with Trypsin. Eur J Biochem. 6, 156-&. (DOI 10.1111/j.1432-1033.1968.tb00433.x)

76 Johnson, L. P., Magee, D. F. 1965 Inhibition of Gastric Motility by a Commercial Duodenal Mucosal Extract Containing Cholecystokinin and Pancreozymin. Nature. 207, 1401-&. (DOI 10.1038/2071401a0)

77 Cooke, A. R. 1967 Effect of Pancreozymin Preparations on Gastric Secretion. Nature. 214, 729-&. (DOI 10.1038/214729a0)

78 Jorpes, E., Mutt, V. 1966 Cholecystokinin and pancreozymin, one single hormone? Acta Physiol Scand. 66, 196–202. (10.1111/j.1748-1716.1966.tb03185.x)

79 Beinfeld, M. C. 2006 CHAPTER 99 - CCK/Gastrin A2 - Kastin, Abba J. In Handbook of Biologically Active Peptides. (ed.^eds. pp. 715–720. Burlington: Academic Press.

80 Menschaert, G., Vandekerckhove, T. T., Baggerman, G., Landuyt, B., Sweedler, J. V., Schoofs, L., Luyten, W., Van Criekinge, W. 2010 A hybrid, de novo based, genome-wide database search approach applied to the sea urchin neuropeptidome. J Proteome Res. 9, 990–996. (10.1021/pr900885k)

81 Cameron, R. A., Samanta, M., Yuan, A., He, D., Davidson, E. 2009 SpBase: the sea urchin genome database and web site. Nucleic Acids Res. 37, D750–754. (10.1093/nar/gkn887)

82 Yu, N., Nachman, R. J., Smagghe, G. 2013 Characterization of sulfakinin and sulfakinin receptor and their roles in food intake in the red flour beetle Tribolium castaneum. Gen Comp Endocrinol. 188, 196–203. (10.1016/j.ygcen.2013.03.006)

83 Brazeau, P., Vale, W., Burgus, R., Ling, N., Butcher, M., Rivier, J., Guillemin, R. 1973 Hypothalamic Polypeptide That Inhibits the Secretion of Immunoreactive Pituitary Growth Hormone. Science. 179, 77.

84 Epelbaum, J. 1986 Somatostatin in the central nervous system: physiology and pathological modifications. Prog Neurobiol. 27, 63–100.

85 Viollet, C., Lepousez, G., Loudes, C., Videau, C., Simon, A., Epelbaum, J. 2008 Somatostatinergic systems in brain: networks and functions. Mol Cell Endocrinol. 286, 75–87. (10.1016/j.mce.2007.09.007)

86 Schlegel, W., Raptis, S., Harvey, R. F., Oliver, J. M., Pfeiffer, E. F. 1977 Inhibition of cholecystokinin-pancreozymin release by somatostatin. Lancet. 2, 166–168.

87 Hayes, J. R., Johnson, D. G., Koerker, D., Williams, R. H. 1975 Inhibition of gastrin release by somatosatin in vitro. Endocrinology. 96, 1374–1376. (10.1210/endo-96-6-1374)

88 Tostivint, H., Lihrmann, I., Vaudry, H. 2008 New insight into the molecular evolution of the somatostatin family. Mol Cell Endocrinol. 286, 5–17. (10.1016/j.mce.2008.02.029)

89 Veenstra, J. A. 2009 Allatostatin C and its paralog allatostatin double C: the arthropod somatostatins. Insect Biochem Mol Biol. 39, 161–170. (10.1016/j.ibmb.2008.10.014)

90 Veenstra, J. A. 2016 Allatostatins C, double C and triple C, the result of a local gene triplication in an ancestral arthropod. Gen Comp Endocrinol. 230-231, 153–157. (10.1016/j.ygcen.2016.04.013)

91 Li, B., Predel, R., Neupert, S., Hauser, F., Tanaka, Y., Cazzamali, G., Williamson, M., Arakane, Y., Verleyen, P., Schoofs, L., et al. 2008 Genomics, transcriptomics, and peptidomics of neuropeptides and protein hormones in the red flour beetle *Tribolium castaneum*. Genome Res. 18, 113–122. (10.1101/gr.6714008)

92 Stewart, M. J., Favrel, P., Rotgans, B. A., Wang, T., Zhao, M., Sohail, M., O’Connor, W. A., Elizur, A., Henry, J., Cummins, S. F. 2014 Neuropeptides encoded by the genomes of the Akoya pearl oyster Pinctata fucata and Pacific oyster Crassostrea gigas: a bioinformatic and peptidomic survey. BMC Genomics. 15, 840. (10.1186/1471-2164-15-840)

93 Cardoso, J. C., Felix, R. C., Bergqvist, C. A., Larhammar, D. 2014 New insights into the evolution of vertebrate CRH (corticotropin-releasing hormone) and invertebrate DH44 (diuretic hormone 44) receptors in metazoans. Gen Comp Endocrinol. 209, 162–170. (10.1016/j.ygcen.2014.09.004)

94 Lee, H. R., Zandawala, M., Lange, A. B., Orchard, I. 2016 Isolation and characterization of the corticotropin-releasing factor-related diuretic hormone receptor in Rhodnius prolixus. Cell Signal. 28, 1152–1162. (10.1016/j.cellsig.2016.05.020)

95 Chang, M. M., Leeman, S. E. 1970 Isolation of a sialogogic peptide from bovine hypothalamic tissue and its characterization as substance P. J Biol Chem. 245, 4784–4790.

96 Studer, R. O., Trzeciak, A., Lergier, W. 1973 Isolierung und Aminosäuresequenz von Substanz P aus Pferdedarm. Helvetica Chimica Acta. 56, 860–866. (10.1002/hlca.19730560307)

97 Us, V. E., Gaddum, J. H. 1931 An unidentified depressor substance in certain tissue extracts. J Physiol. 72, 74–87.

98 Satake, H., Ogasawara, M., Kawada, T., Masuda, K., Aoyama, M., Minakata, H., Chiba, T., Metoki, H., Satou, Y., Satoh, N. 2004 Tachykinin and tachykinin receptor of an ascidian, *Ciona intestinalis:* evolutionary origin of the vertebrate tachykinin family. J Biol Chem. 279, 53798–53805. (10.1074/jbc.M408161200)

99 Schoofs, L., Holman, G. M., Hayes, T. K., Nachman, R. J., De Loof, A. 1990 Locustatachykinin I and II, two novel insect neuropeptides with homology to peptides of the vertebrate tachykinin family. FEBS Lett. 261, 397–401.

100 Jiang, H., Lkhagva, A., Daubnerova, I., Chae, H. S., Simo, L., Jung, S. H., Yoon, Y. K., Lee, N. R., Seong, J. Y., Zitnan, D., et al. 2013 Natalisin, a tachykinin-like signaling system, regulates sexual activity and fecundity in insects. Proc Natl Acad Sci U S A. 110, E3526–3534. (10.1073/pnas.1310676110)

101 Kawada, T., Satake, H., Minakata, H., Muneoka, Y., Nomoto, K. 1999 Characterization of a novel cDNA sequence encoding invertebrate tachykinin-related peptides isolated from the echiuroid worm, *Urechis unicinctus*. Biochem Biophys Res Commun. 263, 848–852. (10.1006/bbrc.1999.1465)

102 Kanda, A., Iwakoshi-Ukena, E., Takuwa-Kuroda, K., Minakata, H. 2003 Isolation and characterization of novel tachykinins from the posterior salivary gland of the common octopus *Octopus vulgaris*. Peptides. 24, 35–43.

103 Kovac, J. R., Chrones, T., Preiksaitis, H. G., Sims, S. M. 2006 Tachykinin receptor expression and function in human esophageal smooth muscle. J Pharmacol Exp Ther. 318, 513–520. (10.1124/jpet.106.104034)

104 Im, S. H., Takle, K., Jo, J., Babcock, D. T., Ma, Z., Xiang, Y., Galko, M. J. 2015 Tachykinin acts upstream of autocrine Hedgehog signaling during nociceptive sensitization in *Drosophila*. Elife. 4, e10735. (10.7554/eLife.10735)

105 Kahsai, L., Kapan, N., Dircksen, H., Winther, A. M., Nassel, D. R. 2010 Metabolic stress responses in *Drosophila* are modulated by brain neurosecretory cells that produce multiple neuropeptides. PLoS One. 5, e11480. (10.1371/journal.pone.0011480)

106 Nassel, D. R. 1999 Tachykinin-related peptides in invertebrates: a review. Peptides. 20, 141–158.

107 Lee, J. H., Miele, M. E., Hicks, D. J., Phillips, K. K., Trent, J. M., Weissman, B. E., Welch, D. R. 1996 KiSS-1, a novel human malignant melanoma metastasis-suppressor gene. J Natl Cancer Inst. 88, 1731–1737.

108 Messager, S., Chatzidaki, E. E., Ma, D., Hendrick, A. G., Zahn, D., Dixon, J., Thresher, R. R., Malinge, I., Lomet, D., Carlton, M. B., et al. 2005 Kisspeptin directly stimulates gonadotropin-releasing hormone release via G protein-coupled receptor 54. Proc Natl Acad Sci U S A. 102, 1761–1766. (10.1073/pnas.0409330102)

109 Copp, D. H., Cheney, B. 1962 Calcitonin-a hormone from the parathyroid which lowers the calcium-level of the blood. Nature. 193, 381–382.

110 Potts, J. T., Jr., Niall, H. D., Keutmann, H. T., Brewer, H. B., Jr., Deftos, L. J. 1968 The amino acid sequence of porcine thyrocalcitonin. Proc Natl Acad Sci U S A. 59, 1321–1328.

111 Foster, G. V., Baghdiantz, A., Kumar, M. A., Slack, E., Soliman, H. A., Macintyre, I. 1964 Thyroid Origin of Calcitonin. Nature. 202, 1303–1305.

112 Friedman, J., Raisz, L. G. 1965 Thyrocalcitonin: inhibitor of bone resorption in tissue culture. Science. 150, 1465–1467.

113 Wendelaar Bonga, S. E., Pang, P. K. 1991 Control of calcium regulating hormones in the vertebrates: parathyroid hormone, calcitonin, prolactin, and stanniocalcin. Int Rev Cytol. 128, 139–213.

114 Furuya, K., Milchak, R. J., Schegg, K. M., Zhang, J., Tobe, S. S., Coast, G. M., Schooley, D. A. 2000 Cockroach diuretic hormones: characterization of a calcitonin-like peptide in insects. Proc Natl Acad Sci U S A. 97, 6469–6474.

115 Veenstra, J. A. 2014 The contribution of the genomes of a termite and a locust to our understanding of insect neuropeptides and neurohormones. Front Physiol. 5, 454. (10.3389/fphys.2014.00454)

116 Sekiguchi, T., Kuwasako, K., Ogasawara, M., Takahashi, H., Matsubara, S., Osugi, T., Muramatsu, I., Sasayama, Y., Suzuki, N., Satake, H. 2016 Evidence for Conservation of the Calcitonin Superfamily and Activity-regulating Mechanisms in the Basal Chordate *Branchiostoma floridae*: INSIGHTS INTO THE MOLECULAR AND FUNCTIONAL EVOLUTION IN CHORDATES. J Biol Chem. 291, 2345–2356. (10.1074/jbc.M115.664003)

117 Amara, S. G., Jonas, V., Rosenfeld, M. G., Ong, E. S., Evans, R. M. 1982 Alternative RNA processing in calcitonin gene expression generates mRNAs encoding different polypeptide products. Nature. 298, 240–244.

118 McLatchie, L. M., Fraser, N. J., Main, M. J., Wise, A., Brown, J., Thompson, N., Solari, R., Lee, M. G., Foord, S. M. 1998 RAMPs regulate the transport and ligand specificity of the calcitonin-receptor-like receptor. Nature. 393, 333–339. (10.1038/30666)

119 Evans, B. N., Rosenblatt, M. I., Mnayer, L. O., Oliver, K. R., Dickerson, I. M. 2000 CGRP-RCP, a novel protein required for signal transduction at calcitonin gene-related peptide and adrenomedullin receptors. J Biol Chem. 275, 31438–31443. (10.1074/jbc.M005604200)

120 Zandawala, M., Paluzzi, J. P., Orchard, I. 2011 Isolation and characterization of the cDNA encoding DH(31) in the kissing bug, Rhodnius prolixus. Mol Cell Endocrinol. 331, 79–88. (10.1016/j.mce.2010.08.012)

121 Zandawala, M. 2012 Calcitonin-like diuretic hormones in insects. Insect Biochem Mol Biol. 42, 816–825. (10.1016/j.ibmb.2012.06.006)

122 Nakabayashi, K., Matsumi, H., Bhalla, A., Bae, J., Mosselman, S., Hsu, S. Y., Hsueh, A. J. 2002 Thyrostimulin, a heterodimer of two new human glycoprotein hormone subunits, activates the thyroid-stimulating hormone receptor. J Clin Invest. 109, 1445–1452. (10.1172/JCI14340)

123 Roch, G. J., Sherwood, N. M. 2014 Glycoprotein Hormones and Their Receptors Emerged at the Origin of Metazoans. Genome Biol Evol. 6, 1466–1479. (10.1093/gbe/evu118)

124 Sudo, S., Kuwabara, Y., Park, J. I., Hsu, S. Y., Hsueh, A. J. 2005 Heterodimeric fly glycoprotein hormone-alpha2 (GPA2) and glycoprotein hormone-beta5 (GPB5) activate fly leucine-rich repeat-containing G protein-coupled receptor-1 (DLGR1) and stimulation of human thyrotropin receptors by chimeric fly GPA2 and human GPB5. Endocrinology. 146, 3596–3604. (10.1210/en.2005-0317)

125 Rocco, D. A., Paluzzi, J. P. 2016 Functional role of the heterodimeric glycoprotein hormone, GPA2/GPB5, and its receptor, LGR1: An invertebrate perspective. Gen Comp Endocrinol. 234, 20–27. (10.1016/j.ygcen.2015.12.011)

126 Cahoreau, C., Klett, D., Combarnous, Y. 2015 Structure function relationships of glycoprotein hormones and their subunits’ ancestors. Front Endocrinol. 6, (ARTN 26 10.3389/fendo.2015.00026)

127 Verleyen, P., Huybrechts, J., Baggerman, G., Van Lommel, A., De Loof, A., Schoofs, L. 2004 SIFamide is a highly conserved neuropeptide: a comparative study in different insect species. Biochem Biophys Res Commun. 320, 334–341. (10.1016/j.bbrc.2004.05.173)

128 Yasuda, A., Yasuda-Kamatani, Y., Nozaki, M., Nakajima, T. 2004 Identification of GYRKPPFNGSIFamide (crustacean-SIFamide) in the crayfish *Procambarus clarkii* by topological mass spectrometry analysis. Gen Comp Endocrinol. 135, 391–400.

129 Verleyen, P., Huybrechts, J., Schoofs, L. 2009 SIFamide illustrates the rapid evolution in Arthropod neuropeptide research. Gen Comp Endocrinol. 162, 27–35. (10.1016/j.ygcen.2008.10.020)

130 Wegener, C., Gorbashov, A. 2008 Molecular evolution of neuropeptides in the genus *Drosophila*. Genome Biol. 9, R131. (10.1186/gb-2008-9-8-r131)

131 Brezina, V., Bank, B., Cropper, E. C., Rosen, S., Vilim, F. S., Kupfermann, I., Weiss, K. R. 1995 Nine members of the myomodulin family of peptide cotransmitters at the B16-ARC neuromuscular junction of *Aplysia*. J Neurophysiol. 74, 54–72.

132 Hewes, R. S., Snowdeal, E. C., 3rd, Saitoe, M., Taghert, P. H. 1998 Functional redundancy of FMRFamide-related peptides at the *Drosophila* larval neuromuscular junction. J Neurosci. 18, 7138–7151.

133 Fujisawa, Y., Furukawa, Y., Ohta, S., Ellis, T. A., Dembrow, N. C., Li, L., Floyd, P. D., Sweedler, J. V., Minakata, H., Nakamaru, K., et al. 1999 The *Aplysia* Mytilus inhibitory peptide-related peptides: Identification, cloning, processing, distribution, and action. Journal of Neuroscience. 19, 9618–9634.

134 Lange, A. B., Bendena, W. G., Tobe, S. S. 1995 The Effect of the 13 Dip-Allatostatins on Myogenic and Induced Contractions of the Cockroach (*Diploptera punctata*) Hindgut. J Insect Physiol. 41, 581–588. (Doi 10.1016/0022-1910(95)00008-I)

135 Criscuolo, A., Gribaldo, S. 2010 BMGE (Block Mapping and Gathering with Entropy): a new software for selection of phylogenetic informative regions from multiple sequence alignments. BMC evolutionary biology. 10, 1.

